# Multiscale genetic architecture of donor-recipient differences reveals intronic LIMS1 locus mismatches associated with long-term renal transplant survival

**DOI:** 10.1101/2022.12.21.521481

**Authors:** Zeguo Sun, Zhongyang Zhang, Khadija Banu, Ian Gibson, Robert Colvin, Zhengzi Yi, Weijia Zhang, Bony De Kumar, John Pell, Arjang Djamali, Lorenzo Gallon, Philip O’Connell, Cijiang He, Jordan Pober, Peter Heeger, Madhav C Menon

**Author notes:** Equal contribution (Zeguo Sun, Zhongyang Zhang).

## Abstract

**Background:** Long-term kidney allograft survival remains suboptimal. Emerging evidence indicates donor-recipient (D-R) mismatches outside of human leukocyte antigens (HLA) contribute to graft survival but mechanisms remain unclear, specifically for those mismatches within intronic regions.

**Methods:** We analyzed genome-wide SNP data of D-R pairs from two well-phenotyped kidney transplant cohorts (median follow-up ~1800 days), Genomics of Chronic Allograft Rejection (GoCAR; n=385) and Clinical Trials in Organ Transplantation 1/17 (CTOT1/17; n=146), quantifying genetic mismatches outside of HLA for every D-R pair at variant, gene, and genome-wide scales.

**Results:** Unbiased genome-wide screen of GoCAR D-R pairs uncovered the LIMS1 locus as the topranked candidate where D-R mismatches associated with death censored graft loss (DCGL). Independent of HLA, a previously unreported relationship between mismatches at a highly linked, intronic haplotype of 30 SNPs was seen as associated with DCGL, with confirmatory association in intra-ancestry D-Rs. Validation testing within the CTOT-01/17 showed similar associations with DCGL. Haplotype D-R mismatches showed a dosage effect, and the introduction of minor alleles in the donor to major allele-carrying recipients showed a greater risk of DCGL. Both the new LIMS1 haplotype and the previously reported LIMS1 SNP rs893403 are expression quantitative trait loci (eQTL) for the gene GCC2 in recipient immune cells, without alterations in GCC2 or LIMS1 coding sequences. Transcriptome enrichment analyses performed on whole blood and within multiple T cell subsets demonstrated significant associations of GCC2 gene, and of either allelic locus, with regulation of TGF-beta-SMAD signaling, implying a role in Treg function and association with rejection.

**Conclusions:** Our analysis unravels intronic non-HLA SNP mismatches within LIMS1 that do not induce protein sequence variation but associate with DCGL. By acting as cis-eQTLs for GCC2 expression, these SNPs modulates TGF-beta signaling and T cell function, associating with immune events and graft outcomes. The findings have clinical implications for risk assessment and individualized therapy in kidney transplant recipients.

## Introduction

In renal transplantation, short-term allograft outcomes including acute rejection have significantly improved, without a proportionate improvement in long-term allograft outcomes^1^. While distinct etiologies are identifiable in half of all allograft loss cases, allograft fibrosis or interstitial fibrosis and tubular atrophy (IF/TA) of unclear etiology account for 30%–40% of cases^2,3^. A role for anti-donor alloimmunity is directly implicated in all cases of allograft failure from rejection, but is also reported in IF/TA (without rejection) by transcriptomic data^4^. Thus, a majority of allograft loss is related to immune injury.

Alloimmune responses leading to allograft damage are triggered by recipient T cells directly interacting with mismatched, polymorphic donor HLA molecules and or by indirectly recognizing donor-derived peptides processed and presented by recipient HLA on recipient antigen presenting cells (APCs). These cognate interactions are driven by DNA sequence mismatch between HLA regions of donor and recipient on chromosome 6. Non-HLA loci that contribute to allograft rejection can alter protein sequences which if mismatched between donor and recipient, can trigger activation of pathogenic, donor-reactive T cells in donor-recipient (D-R) pairs that share HLA, and have been traditionally labeled as “minor” histocompatibility (mH) antigens. Our group among others have employed genome-wide screening to advance this field by demonstrating that such D-R genetic differences can impact allograft survival either via rejection^5–7^ or via allograft IF/TA^8^ independent of HLA matching. Efforts to unravel specific non-HLA gene loci wherein D-R mismatches disproportionately contribute to transplant outcomes, and understanding mechanisms thereof, have clinical applications for allograft risk stratification, surveillance, or even therapeutics. One example is the reported role of mismatches at rs893403 within the LIMS1 gene locus, a copy number variant (CNV)-tagging, intronic single nucleotide polymorphism (SNP), that was associated with rejection episodes in 3 independent cohorts^9^. An increased risk of rejection was observed only when the A allele at this locus was introduced by the donor into G allele-carrying recipients, implying the “directionality” of this allele mismatch.

To provide a more in depth understanding, herein we undertook a systematic and unbiased examination of genome-wide SNP array data from D-R pairs in two prospective kidney transplant cohorts (GoCAR and CTOT01/17)^10,11^ with the goal of identifying mismatches within non-HLA loci that associate with longterm death-censored graft loss (DCGL). After confirming that global D-R genetic differences resulting from SNP mismatches associate with DCGL, we systematically scanned mismatches across all annotated gene loci, to identify individual gene-level mismatches that significantly associated with increased risk of graft loss, independent of HLA. This screening confirmed LIMS1 as a top-ranked gene locus associated with DCGL, independent of genome-wide mismatches. When we further screened SNP-wise D-R mismatches, we identified 30 SNPs in high linkage disequilibrium (LD), distinct from the previously reported rs893403, which are significantly associated with increased risk of graft loss and demonstrated “directionality” when grouped as haplotype alleles. Our analysis of multiple transcriptomic data sets showed a role for both rs893403 and these linked SNPs as cis-expression quantitative trait loci (cis-eQTL) for GCC2 or Golgin-185, a gene adjacent to LIMS1, that promotes TGF-beta signaling in peripheral blood lymphocytes by regulating trafficking of mannose-6-phosphate receptors.

## Results

### Description of study cohorts and D-R mismatches

We employed the subsets of kidney transplant donor-recipient (D-R) pairs with genome-wide genotype information from their parent studies, GoCAR^11^ and CTOT01/17^10,12^, respectively as our discovery and validation cohorts in the current study. Demographic and clinicopathologic characteristics of the D-R pairs in the GoCAR (discovery) and CTOT01/17 (validation) cohorts are shown in **Table S1** and reported in detail elsewhere^8,13^.

#### Discovery cohort

Briefly, the parent GoCAR study was a prospective, multicenter study. Enrolled patients had clinical data, longitudinal blood draws, and serial surveillance biopsies collected at 0, 3, 12, and 24 months, detailed in our published studies^8,11,14^. Genome-wide genotype data was generated for 385 D-R pairs from the GoCAR study followed by an imputation based on the reference panel from the 1000 Genomes Project (Methods). In the GoCAR cohort, the median follow-up time was 1,824 days, 194 (50.4%) were live donors (LDs), and 50 (13.0%) DCGL events were observed (see **Table S1** and methods). After excluding variants with low confidence (INFO score < 0.4) or high missing rate (≥ 0.05), with no alternative allele, or in the MHC region, there remained 30,109,467 variants to calculate genome-wide D-R mismatch score; after filtering variants with low mismatch frequency (≤ 0.05), there remained 2,251,582 common variants used for gene-level and variant-level mismatch calculation (**Figure S1A** and **Methods**).

#### Validation cohort

Briefly, the parent CTOT-01 study was a prospective, multicenter, observational study that enrolled 280 adult and pediatric, crossmatch negative kidney transplant candidates. All CTOT-01 recipients were followed up for 2 years post transplantation, while CTOT-17 was designed to observationally collect data on 5-year outcomes among patients in this cohort (**Methods**). Genome-wide genotype data was generated and imputed as well with the 1000 Genomes Project reference panel for 146 D-R pairs from the parent CTOT-01/17 study (**Methods**). In the CTOT cohort, the median follow-up time was 1,825 days, 124 (84.8%) were living donors, and 9 (6.2%) DCGL events were observed (**Table S1 and Methods**). As compared to the GoCAR cohort, the CTOT cohort had younger donors and recipients, fewer rejection and DCGL events, more donors with African American ancestry, and higher HLA mismatch score, while other relevant variables had no significant difference (**Table S1**). With similar quality control steps as GoCAR on the imputed genotype data, a total of 11,091,731 variants remained for genome-wide D-R mismatch score calculation, and 137,777 SNPs were left for the gene-level and variant-level mismatch analyses (**Figure S1B and Methods**).

#### Defining D-R mismatches at different genomic scales

In any given D-R pair in either cohort, we defined three levels of donor-to-recipient mismatches based on imputed genotype data: genome-wide mismatches, gene-level mismatches, and SNP-level mismatches, where the former two were themselves defined based on cumulative SNP-level mismatches (**Figure 1 and Methods**). At the SNP level, a D-R mismatch was defined as the donor carrying one or two alleles that are not present in the recipient. Depending on the number of different “alien” alleles introduced from the donor to the recipient, a SNP-level mismatch was further categorized as single mismatch (one allele introduced), double mismatch (two alleles introduced), and any mismatch (one or two alleles introduced) (**Figure 1**). We identified 1,280,475 ± 335,138 of any mismatches in the 385 GoCAR D-R pairs and 233,365 ± 97,270 in the 146 CTOT D-R pairs at the genome-wide scale. The non-exonic region contributed dominantly to the genome-wide mismatch in both cohorts and the non-synonymous SNPs contributed ~50% of the mismatches in exonic regions (**Table S2**). The differences in the total number of mismatches between the two cohorts are due to different genotyping platforms and imputation methods, but after normalizing with inter-quartile range (IQR), the genome-wide mismatch score is consistent between studies (see results below).

**Figure 1.**
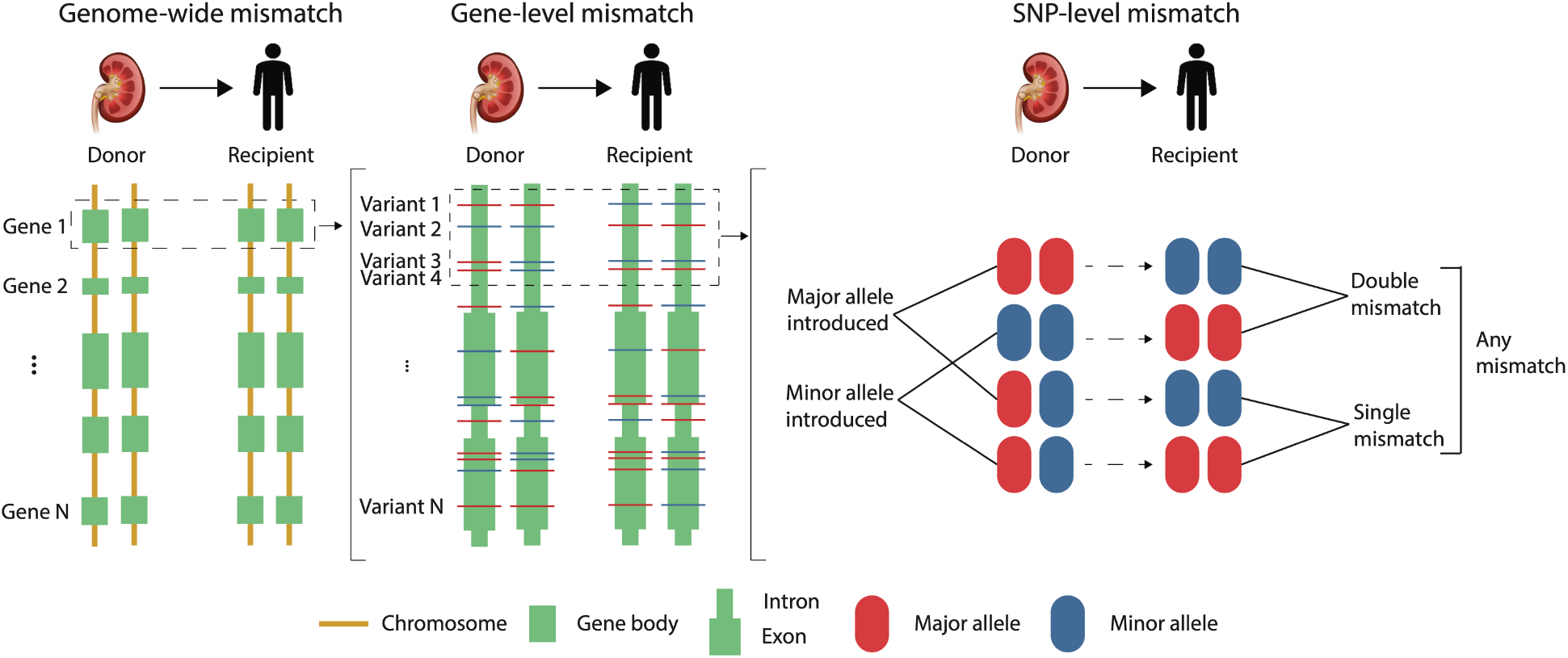
Definition of genetic mismatch between donor-recipient pairs at multiple scales. Genetic D-R mismatches are defined at genome-wide (left), gene (middle), and variant (right) levels, followed by association analysis with transplant outcomes. For a specific variant (e.g., SNP), three different types of mismatches, single mismatch, double mismatch, and any mismatch, are defined as shown in the rightmost part of the diagram.

### Normalized genome-wide non-HLA D-R mismatch score are associated with graft loss

We first calculated the normalized genome-wide non-HLA D-R mismatch score (or simply, the genomewide mismatch score) by summing the SNP-level any mismatch scores (0 for match and 1 for mismatch) at all the imputed, quality-controlled SNPs across the genome (excluding the MHC region) as raw mismatch counts. We normalized the raw counts by their inter-quartile range (IQR) for each D-R pair (**Methods**)^5^. This normalized score was able to capture information from both quantitative measures of genome-wide mismatches that we described in previous work: ^8^ (a) genetic ancestry and (b) proportion of genome-shared identity by descent (pIBD), which are themselves mutually orthogonal.^8^ First, the normalized mismatch score could reflect the relative differences in genetic ancestries of D-R pairs in both GoCAR and CTOT cohorts, where inter-ancestry pairs generally had larger mismatch scores than intra-ancestry pairs (**Figure 2A and C**). Within inter- and intra-ancestry D-R pairs, the distribution of the scores is also consistent with existing knowledge about the relative distance between different major ancestral populations^8^. Second, independent of genetic ancestry in both cohorts, the normalized mismatch score was highly correlated with pIBD, a quantitative measure of kinship between two individuals (**Figure 2B and D**). Hence, we used this variable as a representative quantification of genome-wide D-R mismatches in all association analyses with DCGL. In the discovery cohort, using univariate and multivariable Cox regression (adjusting HLA mismatch score, induction therapy, and donor status), increased genome-wide mismatch scores were associated with DCGL (HR = 1.46, P < 0.001 and HR = 1.25, P = 0.04, respectively) (**Figure 2E**). The association signal in univariate model was validated in the CTOT cohort, while we observed similar HR in multivariable model albeit less significant with wider confidence interval due to lower sample size and event rate (**Figure 2E**). As sensitivity analyses, we evaluated European-to-European (E-to-E) D-R pairs and identified similar association signals (**Figure S2**).

**Figure 2.**
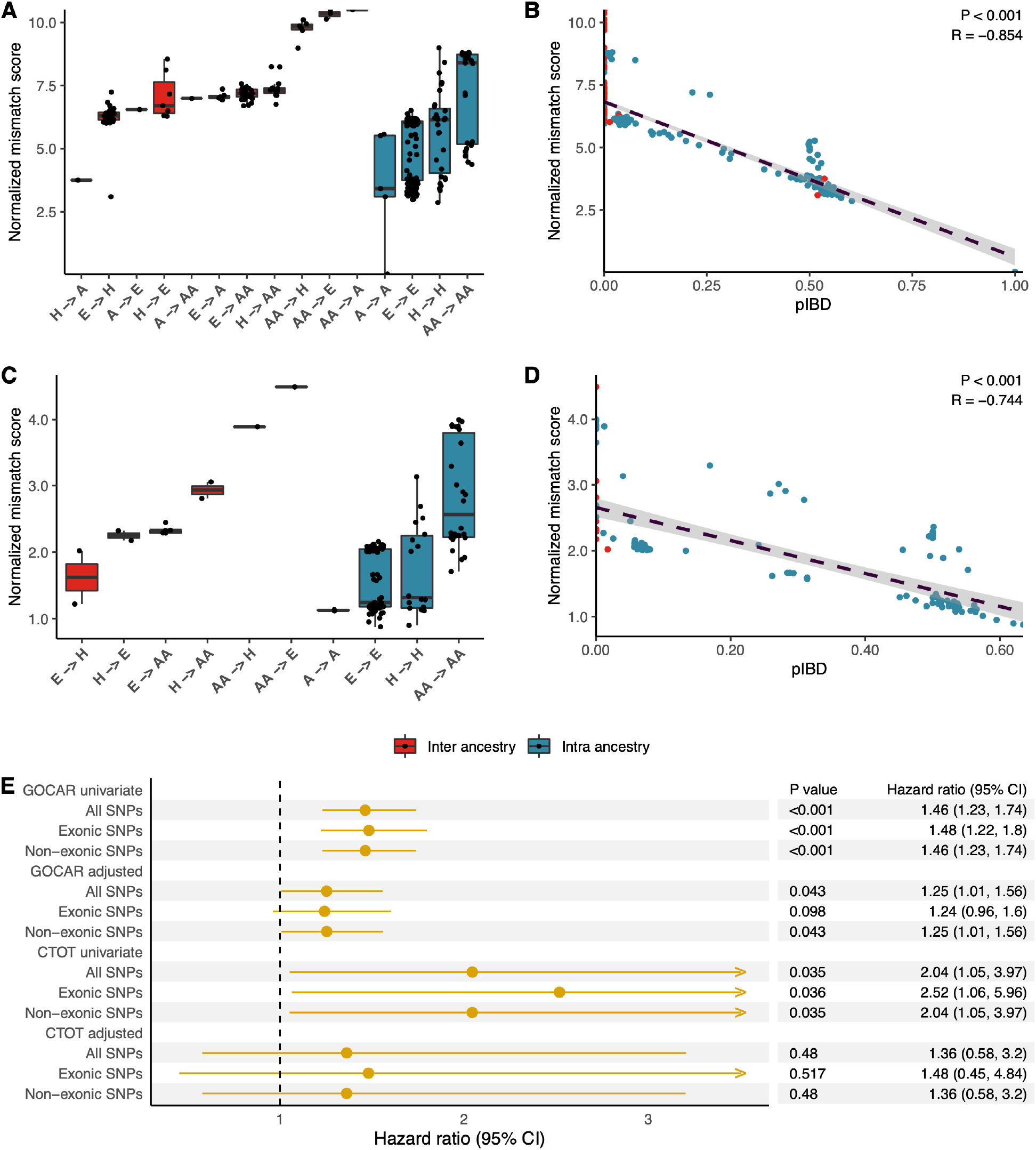
Genome-wide D-R mismatch score was associated with graft loss. Genome-wide mismatch score between donor-recipient (D-R) pairs was calculated at all imputed SNPs with high confidence after QC, and normalized by inter-quartile range (IQR). The distribution of normalized genome-wide mismatch scores is shown for D-R pairs stratified by different combinations of D-R genetic ancestries in GOCAR (**A**) and CTOT (**C**), with inter-ancestry D-R pairs in red and intra-ancestry in blue. Genome-wide D-R mismatch score is highly correlated with inter-personal relatedness, reflected as the proportion of identity-by-decent (pIBD) in GOCAR (**B**) and CTOT (**D**). (**E**) Forest plots show the association with DCGL of genome-wide D-R mismatch scores calculated at all imputed SNPs as well as the SNPs within exonic and non-exonic regions. The association analyses were performed using univariate and multivariable Cox regression models adjusted for HLA mismatches, induction therapy, and donor status for both the GOCAR and CTOT cohorts.

Within our cohorts, we also aimed to evaluate genome-wide mismatches defined within transmembrane non-synonymous SNPs vs. other genomic regions and tested their reported association with DCGL (**Methods**)^5^. We observed high correlations between these different components of the normalized genome-wide mismatch score (**Figure S3**). Hence, the specific role of transmembrane non-synonymous SNP mismatches and their impact on graft survival could not be dissected in our datasets.

### Screening of gene-level mismatches unraveled mismatches at LIMS1 locus associated with graft loss

We next scanned the whole genome in an effort to pinpoint specific non-HLA gene loci at which D-R mismatches are associated with graft loss. To achieve this, we derived gene-level mismatch scores by summing over SNP-level mismatch scores for SNPs mapped to each gene region (**Methods**). To avoid identification of rare-variant based mismatches while enriching for important gene-level signals, we only considered SNPs with relatively frequent occurrence of mismatches (≥ 5% D-R pairs in our study cohorts) (**Figure S1 and Methods**). We then screened gene specific mismatch scores for the association with DCGL, each using a multivariable Cox regression adjusting genome-wide mismatch score, HLA mismatch score, and clinical covariates respectively (**Figure 3A**). We considered any mismatch and double mismatch for all and E-to-E D-R pairs, resulting in four different settings for analyses. In GoCAR, among 20,141 genes with derived mismatch score, we identified that gene-level mismatch score at LIMS1 locus was robustly associated with DCGL in all four scenarios, independent of genome-wide mismatch, highlighting the important role of this non-HLA locus in graft outcomes (**Figure 3B and C, Figure S4B and C, Table S3**). The association signal can also be validated in the CTOT cohort (**Table S4**). As sensitivity analysis, a less stringent threshold (nominal p ≤ 0.05) led to a list of 23 candidate genes (including LIMS1) whose gene-level mismatch scores were associated with DCGL (**Table S3 and Figure S4A**). We summarized mismatches at the 23 gene loci into a single mismatch score by accumulating gene-wise mismatch score, and evaluated its association with DCGL. In adjusted Cox models, the 23-gene mismatch score was significantly associated with DCGL, while the association of the remaining genome mismatches with DCGL was markedly attenuated, highlighting the disproportionate relevance of mismatches in these 23 non-HLA genes (**Figure S5**). Notably, among the 23 genes, GCC2 and GCC2-AS1, located on Chromosome 2q12, are in proximity to LIMS1. Hence these analyses implicated a role for mismatches in the non-HLA chromosomal region surrounding the LIMS1 gene.

**Figure 3.**
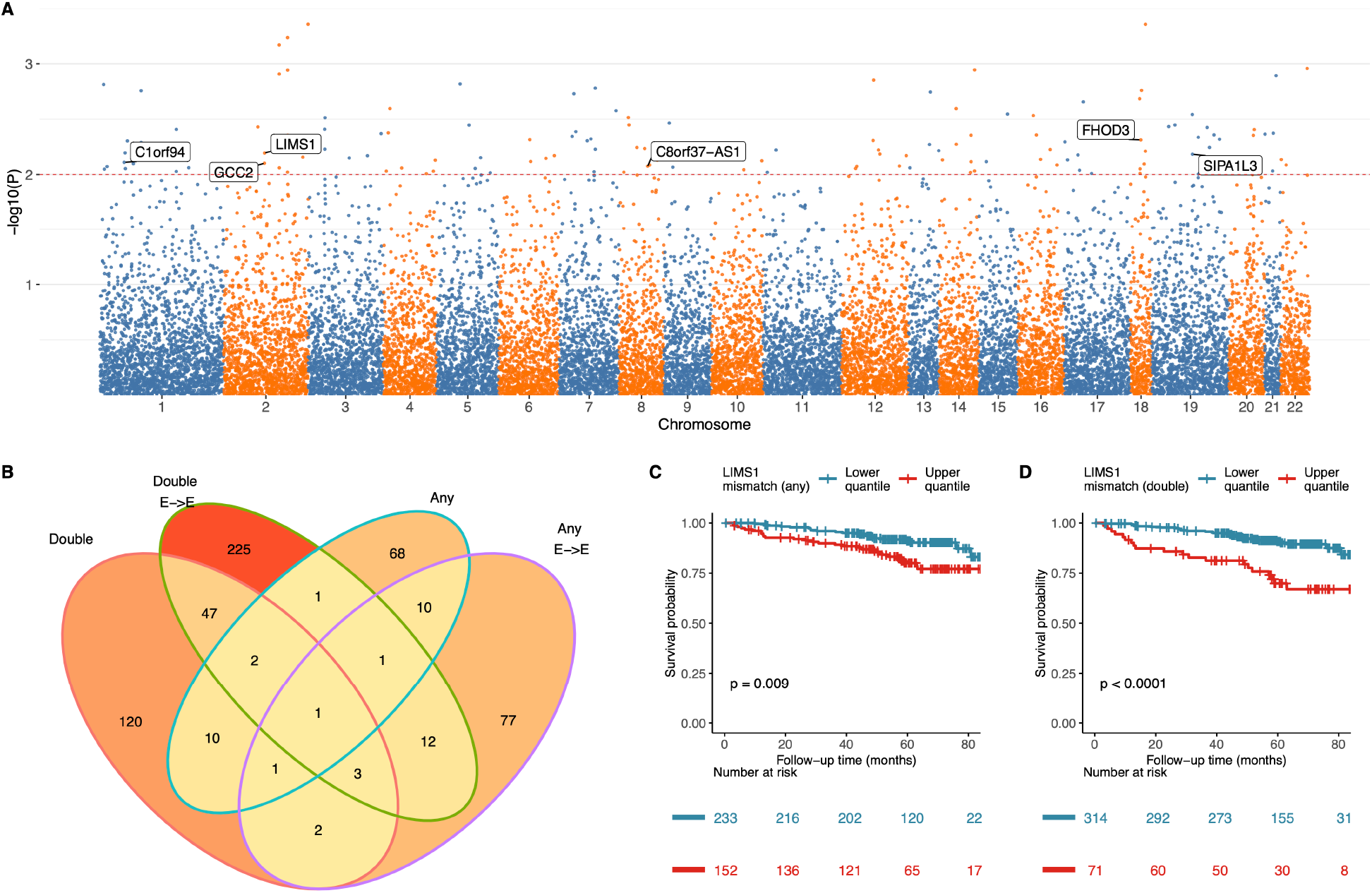
Genome-wide screening of D-R mismatch at gene-level revealed the association of mismatch score at LIMS1 locus with graft loss. (**A**) A Manhattan plot shows the p-values of the association of gene-level any mismatch scores with DCGL in GOCAR. Top candidate gene loci from four different analyses shown in **(B)** are highlighted. (**B**) Venn diagram shows the number of genes identified with mismatch score at LIMS1 locus in significant association with DCGL (nominal p ≤ 0.01) from four different analyses: double mismatch or any mismatch (definition in Figure 1 and Methods) for the whole GOCAR cohort or the subset of European-to-European (E-to-E) D-R pairs. Kaplan-Meier plots show the graft survival curves for equally dichotomized groups of mismatch scores at LIMS1 locus, where mismatch scores were defined as “any mismatch” in (**C**) and “double mismatch” in (**D**). P-values were derived from log-rank tests in comparison of upper quantile versus lower quantile.

### Evaluation of SNP-level mismatches in LIMS1 revealed novel variants associated with DCGL

As the top candidate gene locus in our study, LIMS1 was also implicated in previous studies^9,15^. The mismatch at intronic SNP rs893403 within LIMS1, defined by presence of homozygosity in the CNV-tagging minor allele in the recipient and presence of major allele (either heterozygous or homozygous) in the donor kidney, was reported to be associated with increased risk of rejection^9^. In GoCAR, we observed mismatch at rs893403 associated with increased risk of DCGL in all and E-to-E D-R pairs (**Figure 4A and B**).

**Figure 4.**
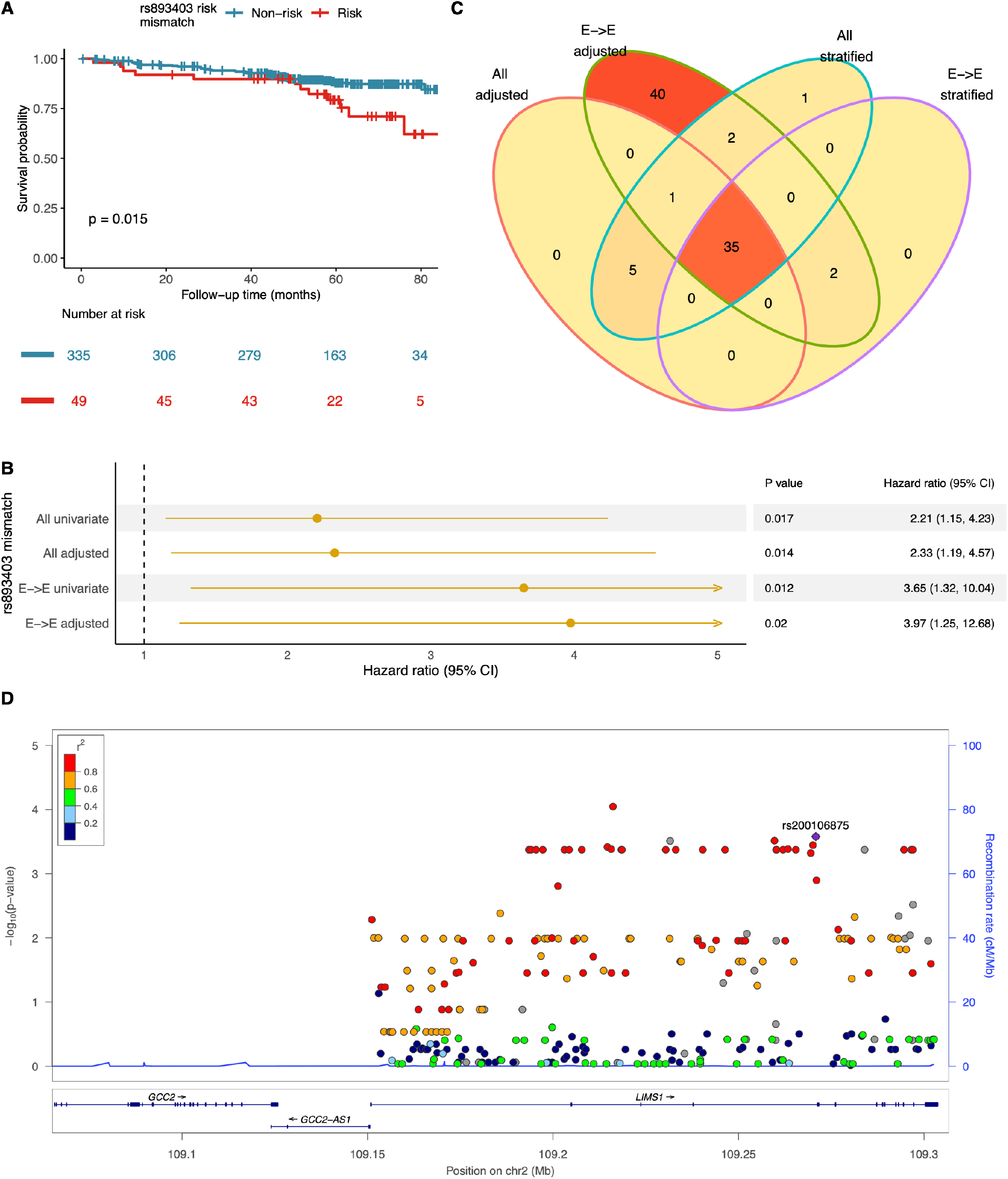
Variant-level mismatches at LIMS1 locus were associated with death-censored graft loss. (**A**) Kaplan-Meier plot shows the SNP rs893403 mismatch was associated with DCGL. P-value was derived from log-rank test in comparison of mismatch versus non-mismatch group. (**B**) Forrest plot for the association of the rs893403 mismatch with DCGL in all and European-to-European D-R pairs with univariate and multivariable model adjusted genome-wide mismatch score, HLA mismatch score, donor status, and induction therapy. (**C**) Venn diagram shows the number of top candidate SNPs (other than rs893403) associated with DCGL (nominal p ≤ 0.05) within the LIMS1 gene region in Cox regression analyses adjusted by rs893403 mismatch or within the rs893403 non-risk stratum for all and E-to-E D-R pairs. (**D**) LocusZoom plot shows the association p-values (on minus log10 scale) of variants with high frequency of any mismatch (≥ 5% D-R pairs) within LIMS1 locus in adjusted analysis in all D-R pairs. The linkage disequilibrium metric R^2^ was calculated for SNPs surrounding one of the top candidate SNPs, rs200106875, within the LIMS1 gene region.

We then evaluated SNP-level D-R mismatches (any mismatch) for all non-rs893403 LIMS1 SNPs (n = 287 variants after QC; see methods) and their individual associations with DCGL. To investigate association signals independent of rs893403, we screened any mismatch of each LIMS1 SNP using (a) Cox models adjusted by rs893403 risk mismatches (n = 385), and (b) stratified Cox model by excluding D-R pairs with rs893403 risk mismatches, i.e., within the rs893403 non-risk stratum (n = 335). (**Figure 4C**). The SNP-wise screening was carried out in all and E-to-E D-R pairs, resulting in four analyses for each SNP. **Table S5** shows the top ranked (by p-value) 30 individual SNP-mismatches, which were identified as significantly associated with DCGL in all the four scenarios. Notably, all the 30 SNPs identified are located in LIMS1 intronic regions. Using the linkage disequilibrium (LD) data derived from all the major continental populations of 1000 Genomes Project^16^, we observed that these intronic SNPs are in high LD with each other (R^2^ > 0.99), but are not linked with rs893403 (R^2^<0.5) (**Figure S6A**), while the latter itself serves as a tagging SNP of a different haplotype within the LIMS1 locus (**Figure S6B**). Hence, we accounted for mismatches within these linked SNPs as a single block (or haplotype) distinct from rs893403 (**Figure 4D** and **Figure S6**). In the CTOT (validation) cohort, we successfully reidentified 28 of the 30 haplotype SNPs discovered in GoCAR. Here too, D-R mismatch within this haplotype was associated with increased risk of DCGL in univariate and adjusted multivariable survival analyses (**Figure S7 and Table S6**).

In summary, we confirmed and extended prior data regarding the LIMS1 locus implicated in kidney transplant rejection, and specifically the D-R mismatch at the CNV-tagging SNP rs893403, and identified its association with DCGL. We also discovered that novel and independent mismatch within the LIMS1 locus (surrogated by a 30-SNP haplotype) was associated with DCGL, overall showing the importance of this chromosomal region in allotransplantation.

### Dosage effect and directionality of donor-recipient mismatch at the LIMS1 haplotype

Using Cox regression, our analysis showed that the number of haplotype allele mismatches were associated with increased risk of DCGL in all and E-to-E D-R pairs in GoCAR, suggesting a dosage effect for D-R mismatches at this haplotype (**Figure 5A and B and Table 1**)^9^. In GoCAR cohort, we showed that the association of rs893403 mismatch with DCGL had directionality: only donor A allele introduced into recipients with homozygous G allele (linked to the deletion) increased the risk of graft loss, consistent with existing results on the directionality of the association with acute rejection^9^. We next explored the directionality of the effect on DCGL of the newly identified haplotype mismatch: donor kidney introduced minor allele of the haplotype to a mismatched recipient carrying homozygous major allele vs. vice versa. To minimize confounding from rs893403 mismatch, we set apart the D-R pairs with rs893403 risk mismatch into a single group (regardless the mismatch status of the haplotype). In the whole cohort, and in E-to-E D-R pairs, patients with rs893403 risk mismatch and minor haplotype allele-introducing mismatch had worse DCGL outcomes (**Figure 5C and D and Table 2**). In the E-to-E D-R pairs, the major haplotype allele-introducing mismatch was not significantly associated with DCGL. Overall, our analyses revealed an observable dosage effect on DCGL from LIMS1 haplotype mismatch and the directionality of this effect, i.e., worse allograft outcomes observed in D-R pairs with minor-to-major haplotype mismatch than major-to-minor mismatch, even after accounting for the rs893403 risk mismatch.

**Figure 5.**
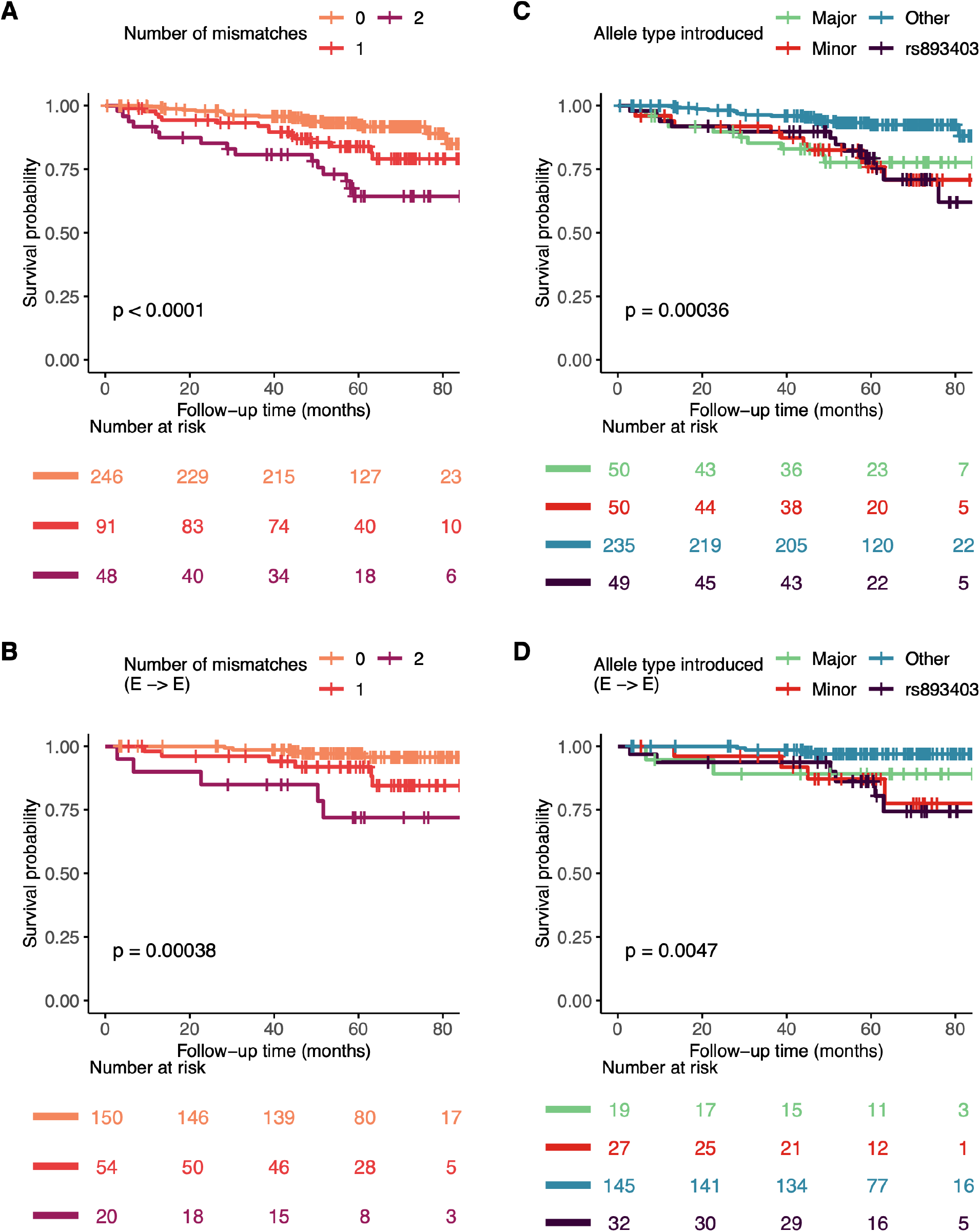
Dosage effect and directionality of the haplotype mismatch associated with DCGL. Kaplan-Meier survival curves of DCGL grouped by the dosage of mismatches (i.e. the number of “alien” allele introduced by the donor) at the haplotype in all (**A**) and European-to-European (E-to-E) D-R pairs (**B**). Kaplan-Meier survival curves of DCGL grouped by the directionality of the mismatches at the haplotype and the rs893403 risk mismatch in all (**C**) and E-to-E D-R pairs (**D**). Major: mismatch derived from major haplotype allele introduced by donor to the recipient with homozygous minor allele and no rs893403 risk mismatch; Minor: mismatch derived from minor haplotype allele introduced by donor to the recipient with homozygous minor allele and no rs893403 risk mismatch; rs893403: mismatch at rs893403 defined as risk allele (A allele) introduced by donor to the recipient carrying G/G genotype; Other: no mismatch at the haplotype and rs893403.

**Table 1.** Dosage effect of the haplotype mismatch in association with DCGL in the GoCAR cohort.

| Variable | HR | 95% CI | P value |
| --- | --- | --- | --- |
| **All D-R pairs (n = 384; 50 [13.0%] graft loss events)** |  |  |  |
| **Dosage of haplotype mismatches (ref: no mismatch)** |  |  |  |
| 1 mismatch | 2.85 | (1.34, 6.05) | **0.01** |
| 2 mismatches | 3.95 | (1.85, 8.46) | **<0.001** |
| Genome-wide mismatch | 1.55 | (1.07, 2.24) | **0.02** |
| Induction (ref: No) | 2.66 | (0.93, 7.60) | 0.07 |
| Donor status (ref: live donor) | 2.05 | (1.05, 4.02) | **0.04** |
| HLA mismatch score | 1.03 | (0.73, 1.46) | 0.87 |
| rs893403 risk mismatch (ref: no) | 1.58 | (0.79, 3.18) | 0.20 |
| **E-to-E D-R pairs (n = 223; 17 [7.6%] graft loss events)** |  |  |  |
| **Dosage of haplotype mismatches (ref: no mismatch)** |  |  |  |
| 1 mismatch | 5.29 | (1.23, 22.79) | **0.03** |
| 2 mismatches | 9.66 | (2.25, 41.51) | **<0.001** |
| Genome-wide mismatches | 2.16 | (0.78, 6.03) | 0.14 |
| Induction (ref: No) | 2.17 | (0.58, 8.09) | 0.25 |
| Donor status (ref: live donor) | 1.08 | (0.36, 3.3) | 0.89 |
| HLA mismatch score | 0.75 | (0.44, 1.28) | 0.29 |
| rs893403 risk mismatch (ref: no) | 1.79 | (0.53, 6.01) | 0.35 |

**Table 2.** The directionality of the haplotype mismatch in association with DCGL in the GoCAR cohort.

| Variable | HR | 95% CI | P value |
| --- | --- | --- | --- |
| **All D-R pairs (n = 384; 50 [13.0%] graft loss events)** |  |  |  |
| **Mismatch group<sup>a</sup> (ref: no mismatch)** |  |  |  |
| rs893403 | 4.52 | (2.04, 10) | **<0.001** |
| Minor | 4.00 | (1.79, 8.93) | **<0.001** |
| Major | 3.75 | (1.61, 8.75) | **<0.001** |
| Genome-wide mismatch | 1.52 | (1.05, 2.18) | **0.03** |
| Induction (ref: No) | 2.7 | (0.94, 7.72) | 0.06 |
| Donor status (ref: LDs) | 2.12 | (1.06, 4.2) | **0.03** |
| HLA-mismatch score | 1.04 | (0.73, 1.47) | 0.84 |
| **E-to-E D-R pairs (n = 223; 17 [7.6%] graft loss events)** |  |  |  |
| **Mismatch group<sup>a</sup> (ref: no mismatch)** |  |  |  |
| rs893403 | 11.09 | (2.57, 47.86) | **<0.001** |
| Minor | 10.15 | (2.36, 43.72) | **<0.001** |
| Major | 6.39 | (0.93, 43.7) | 0.06 |
| Genome-wide mismatches | 2.11 | (0.77, 5.81) | 0.15 |
| Induction (ref: No) | 2.30 | (0.61, 8.68) | 0.22 |
| Donor status (ref: live donors) | 1.24 | (0.37, 4.08) | 0.73 |
| HLA mismatch score | 0.75 | (0.44, 1.28) | 0.29 |

<sup>a</sup>: Definition of the mismatch group. Major: mismatch derived from major haplotype allele introduced by donor to the recipient with homozygous minor allele and no rs893403 risk mismatch; Minor: mismatch derived from minor haplotype allele introduced by donor to the recipient with homozygous minor allele and no rs893403 risk mismatch; rs893403: mismatch at rs893403 defined as risk allele (A allele) introduced by donor to the recipient carrying G/G genotype; Reference: no mismatch at the haplotype and rs893403.

### LIMS1 SNPs are expression quantitative trait loci (eQTLs) for *GCC2* associated with Treg activation and TGF-beta-SMAD signaling

To explore the putative mechanism of the directional mismatches of the SNPs in intronic region of LIMS1 in contributing to DCGL, and based on the previous report of rs893403 as an eQTL in non-glomerular renal tissue (**Figure S8A**)^9^, we evaluated the association between LIMS1 gene expression and the haplotype. To study the *donor* haplotype and gene expression in allograft kidneys, we examined the NephQTL data of published kidney transcriptomes^17^. Similar to rs893403, the minor allele of the haplotype SNPs were associated with increased expression of LIMS1 in tubules (**Figure S8B**).

To simultaneously explore the role of the haplotype alleles in the recipient during the occurrence of a “mismatch”, we evaluated the eQTL function in immune cell transcriptomes from published data^18,19^. Unexpectedly, from multiple datasets, the LIMS1 haplotype is not an eQTL for *LIMS1* in peripheral blood mononuclear cells (PBMCs). Instead, both the LIMS1 haplotype and rs893403 demonstrated significant eQTL function on *GCC2*, the 5’-end neighboring gene to LIMS1 (**Figure 6A and Figure S9**). GCC2 (Golgin 185) is a GRIP-domain containing protein with canonical role in late endosome-to-Golgi trafficking by binding Rab- and Arl1-GTPases^20–22^ and is essential for mannose-6-phosphate receptor (M6PR) recycling^22^. In turn, M6PR proteins have roles in binding and activation of latent TGF-beta by cleavage of latent peptide (promoting TGF-beta signaling)^23,24^, and in insulin-like growth factor-2 (IGF-2) signaling^24–26^.

**Figure 6.**
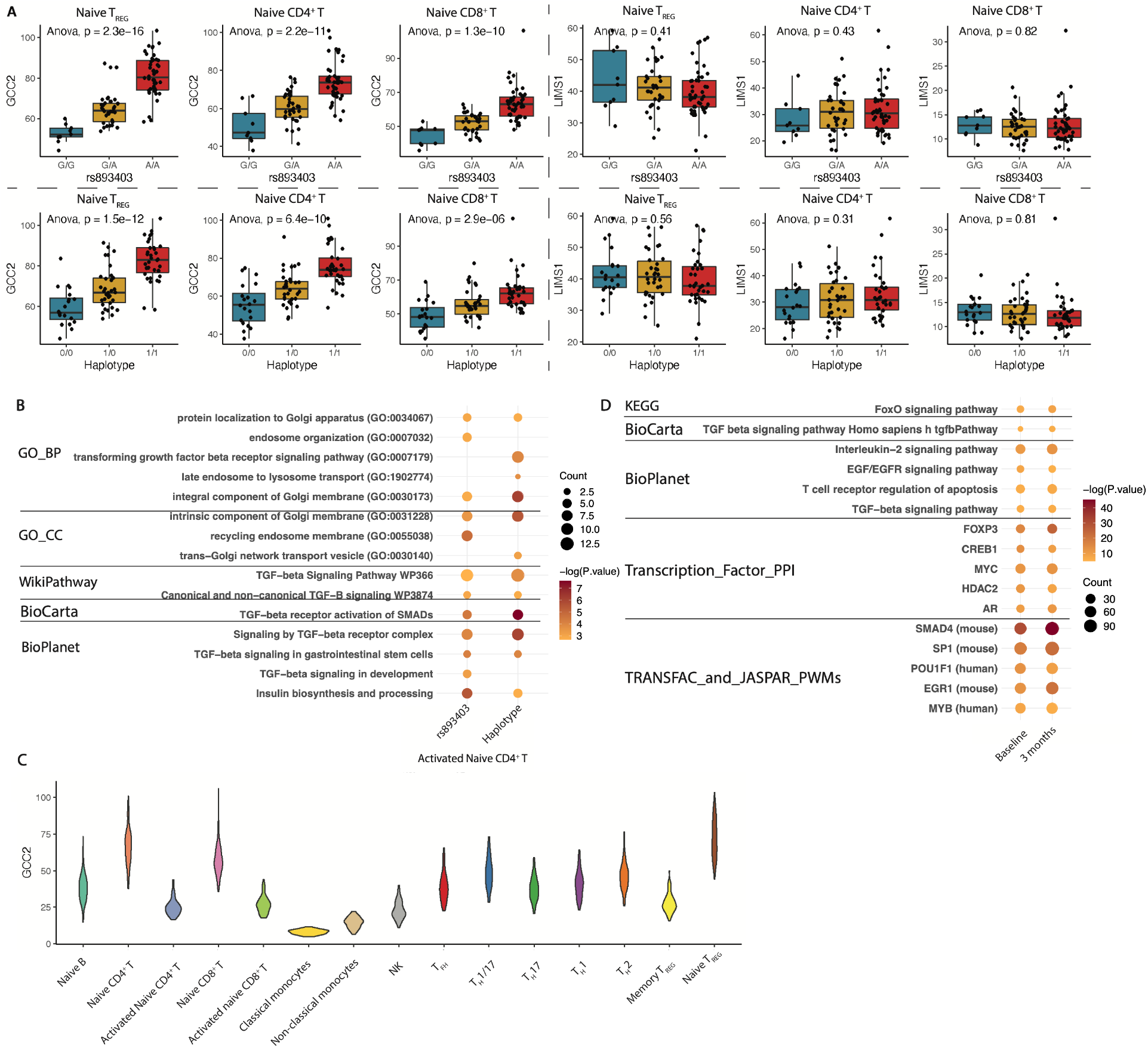
eQTL analysis of rs893403 and the haplotype revealed GCC2 as cis-regulated gene. (**A**) Box plots show the distribution of GCC2 and LIMS1 expression within each genotype group of the haplotype and rs893403 in naive Treg, naive CD4^+^ T cell, and naive CD8^+^ T cell from the DICE cohort. The significance of the association between expression level and the genotype is indicated by the p-value derived from ANOVA. (**B**) Enriched pathways of DEGs (nominal p ≤ 0.05) associated the number of risk alleles of rs893403 and the haplotype in naive CD4^+^ T cell from the DICE cohort. (**C**) The distribution of GCC2 expression, displayed as violin plot, in the 15 different immune cell types from the DICE cohort. (**D**) Enriched pathways and transcription factors of genes positively co-expressed with GCC2 (R ≥ 0.6, p ≤ 0.01) in pre-transplant and 3-month post-transplant patients from the GOCAR cohort. The pathways or transcription factors shown in (C) and (D) are grouped by the source databases.

Using the published DICE (Database of Immune Cell Expression, Expression quantitative trait loci and Epigenomics) cohort^18^, where sorted PBMCs after leukapheresis from healthy controls were genotyped and underwent RNA-seq (**Methods**), we successfully re-identified 22 of the 30 haplotype SNPs, which also showed consistent genotypes across samples. From this data of sorted immune cell populations, **Table S7** shows the rank of *GCC2* among genes across the transcriptome (by Benjamini-Hochberg adjusted P-value) in the eQTL analysis of rs893403 and the haplotype in each of the 15 sorted PBMC subtypes, among which naive Tregs were top-ranked (**Methods**). We also evaluated transcriptome-wide differentially expressed genes (DEGs) based on rs893403 and the LIMS1 haplotype genotype (**Methods**). Consistent with the described role of GCC2, we identified enrichment of endosomal transport, TGF-beta-SMAD signaling, and IGF-signaling in these analyses (**Figure 6B** and **Figure S10**).

We next studied GCC2 expression levels in immune cell subsets of the DICE data. *GCC2* was highly expressed in naive Tregs, naive CD4^+^ and CD8^+^ T cells, and was downregulated upon T cell receptor stimulation (**Figure 6C**). *LIMS1* expression was relatively higher in monocytes (**Figure S11A**). Relative expressions of *GCC2* and *LIMS1* in PBMC subsets were corroborated in our own PBMC single cell sequencing data obtained from end-stage kidney disease patients (**Figure S11B**)^13^.

To explore the function of GCC2 in peripheral blood cells using data from the parental GoCAR study, we performed co-expression analyses using previously obtained whole blood transcriptomes from recipients, both pre-transplantation (GSE112927) and at 3 months post-transplant (GSE120398) (n = 292 and 147 respectively). As shown in **Figure 6D**, GCC2 co-expressed genes (R ≥ 0.6; Adjusted P ≤ 0.05) (n = 571 and 853 genes respectively) showed significant and consistent enrichment of TGF-beta-SMAD signaling pathway terms and FOXP3 transcription factor targets throughout multiple databases (also see **Methods**). GCC2 co-expressed genes (R ≥ 0.6; P ≤ 0.05) in each cell type from the healthy controls of the DICE cohort also showed enrichment of TGF-beta-SMAD signaling pathway terms and FOXP3 in multiple T cell subsets (**Figure S12** and Methods).

Our analysis here implies a regulatory role for the rs893403 and the LIMS1 haplotype on *GCC2* expression in immune cell subsets including Tregs, and further links *GCC2* expression with the described canonical function relating to TGF-beta-SMAD signaling in immune cells.

### In silico analyses demonstrate SNPs in high LD with LIMS1-SNPs and cis-eQTL function for GCC2

Since our above analyses suggested eQTL functions of rs893403 and the LIMS1 haplotype for *GCC2,* we aimed to further prioritize these SNPs by integrating publicly available datasets including Assay for Transposase-Accessible Chromatin with high-throughput sequencing (ATAC-seq) data^27^, ChIP-seq data^28^, DNAse-hypersensitivity^29^, transcription factor binding ^29^, and ENCODE data. Our goal was to identify putative “causal” SNPs in our list of candidate LIMS1 SNPs including rs893403. As shown in **Figure 7,** rs893403 is located in an ATAC-seq peak region specific to immune cells, while there are no peaks detected within the previously described CNV (CNVR915.1) tagged by rs893403. Evaluation of SNPs highly linked (R^2^>0.9) to rs893403 revealed two candidate SNPs located in accessible chromatin areas, rs2460944 (R^2^=0.96) within the transcription start site (TSS) of GCC2 and rs10084199 (R^2^=0.91) in the TSS of LIMS1 respectively. These SNPs showed DNase hypersensitivity, high RegulomeDB scores (**Methods**)^30^, transcription favorable histone modifications, and confirmed highly significant eQTL function in multiple tissues for GCC2 based on GTEX data (**Tables S8-9**). SNP rs2460944 mismatches were found to be associated with DCGL with similar directionality as rs893403 (**Figure S13)**.

**Figure 7.**
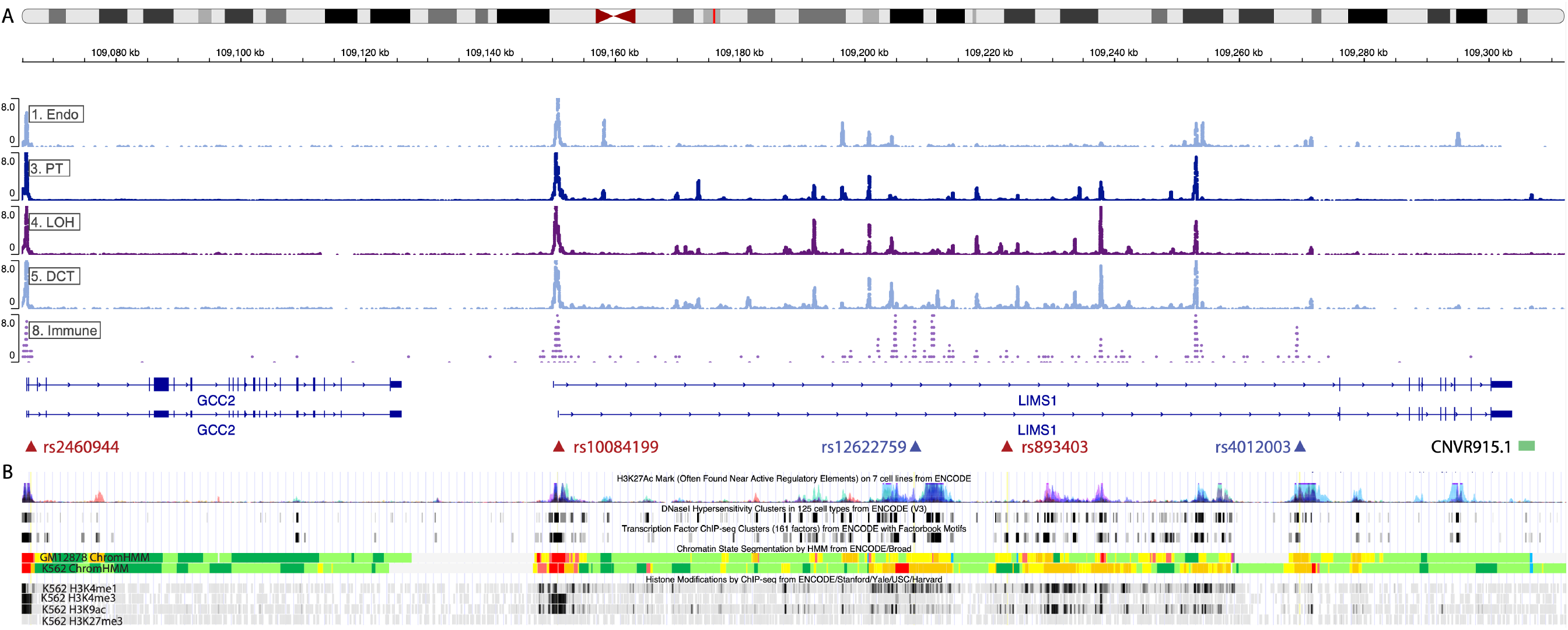
Potential regulatory SNPs in high LD with rs893403 and the haplotype. (**A**) Cell-specific peaks from whole kidney single cell ATAC-seq data^27^. SNP IDs in red are in LD with rs893403, while SNP IDs in blue are in LD with the haplotype. (**B**) Annotation of epigenetic and transcription factor ChIP-seq data in corresponding tracks from the UCSC genome browser.

Among candidate SNPs within the LIMS1 haplotype, rs4012003, and an additional SNP, rs12622759, are located in areas of open chromatin, with further evidence of DNase hypersensitivity, high RegulomeDB scores, favorable histone modifications, and significant eQTL function for GCC2 in multiple cell types. These peak regions appear specific to immune cells according to the scATAC-seq data. These *in silico* analyses suggest putative “causal” SNPs within the LIMS1-GCC2 locus which are tagged by rs893403 or within the LIMS1 haplotype and could regulate GCC2 expression.

## Discussion

The traditional paradigm regarding mismatches between donor and recipient is the development of an adaptive immune response against donor proteins of dissimilar peptide sequence by the recipient’s allorecognition mechanisms. These responses are primarily determined by mismatches between the HLA regions of donor and recipient.^5-8^ Here, a systematic approach screening the whole genome to discover specific non-HLA gene loci revealed LIMS1 D-R mismatches as a top-ranked candidate associated with DCGL. Within the LIMS1 locus, the role and directionality of mismatches at the previously reported intronic SNP rs893403, was extended to long-term DCGL. In addition to rs893403, a novel haplotype of 30 intronic LIMS1 SNPs, almost perfectly linked with each other, was identified where mismatches were independently associated with increased risk of DCGL, again with a demonstrable directionality (minor allele in donors introduced into major allele-carrying recipients). Hence our analyses provide a pipeline for identifying the role of non-HLA gene mismatches relevant to allograft survival.

Most interestingly, neither rs893403 nor the SNPs in the LIMS1 haplotype alter LIMS1 protein sequence; instead, both were identified as cis-eQTLs for an adjacent gene, GCC2, in immune cells with high expression in Tregs. Our subsequent transcriptomic analyses revealed the association of LIMS1 SNPs with TGF-beta-SMAD signaling via GCC2 expression in multiple T cell subsets – including regulatory T cells and naive CD4^+^ and CD8^+^ T cells. These transcriptomic data are consistent with the expected canonical role of GCC2 (or Golgin-185) as a trans-Golgi network protein in the Golgi-to-endosome trafficking of M6PRs. Specifically, Cation-independent M6PR activates TGF-beta by cleaving the latent peptide and acts as a receptor for IGF-2 signaling (**Figure 6B**).

While we validated and extended previous association of the LIMS1 rs893403 mismatch to allograft survival, we could not find significant associations with either ABMR^9^ or TCMR^15^ in our datasets. We note that our study cohorts reported subclinical and clinical rejection episodes up to 2 years by respective central pathology cores, but later episodes were not required to be reported in these datasets. Indeed, we identified an association of minor allele at LIMS1 haplotype, or alternatively A allele at rs893403 (accompanying a high-risk mismatch), with allograft fibrosis score at 12 months (CADI or Ci+Ct in GOCAR; not shown), which we previously reported as associated with DCGL^11^. These data are consistent with the prior role for LIMS1 in fibrosis via facilitating ILK-signaling and incomplete epithelio-mesenchymal transition in tubular cells, another potential mechanism linking donor minor haplotype allele to adverse allograft outcomes^31^. In addition, these data could also be consistent with a role for GCC2 itself in renal tubular cells, with minor haplotype allele-carrying donor allografts associated with increased IF/TA via increased TGF-beta signaling ^32,33^, which we and others have reported as associated with early allograft fibrosis by gene expression profiling. On the other hand, the deletion-tagging allele (G allele) in recipients with a high-risk mismatch (i.e., A allele donors to G/G recipients)^9^ was associated with lower GCC2 levels in T cells and likely affected Treg function, via reduced TGF-beta-SMAD signaling. Indeed, the prior reported detection of anti-LIMS1 antibodies in recipients with a high-risk mismatch could itself reflect impairment of Treg function in G/G recipients, and a break in tolerance. Hence intronic regulatory loci in the context of a “directional” LIMS1 mismatch exert cell-specific effects by modulating gene expression and in turn impact allograft survival. Hence, we shed new light on the association of LIMS1 mismatches with allograft outcomes by identifying these intronic SNPs as eQTLs for GCC2 in T cells, impinging on its expected role in TGF-beta signaling, and revealing an unexpected mechanism outside of the usual paradigm involving non-HLA donor-recipient mismatches.

Our data has potential clinical implications. In combination with the prior reports ^9,15^, our data bring out the important role of LIMS1-locus mismatches and this non-HLA genomic region in renal allotransplantation. Genotyping for these variants to delineate a high-risk mismatch could potentially allow for risk stratification – for personalized optimization of immunosuppression, or early surveillance for IF/TA or rejection (based on the prior work), or for enriching patient enrollment in subsequent clinical trials based on LIMS1 mismatches. Given the organ shortage, we believe LIMS1 mismatches are not likely to be utilized for organ allocation, except in exceptional circumstances (for instance where two living donors are considered for the same recipient). Nonetheless, our identification of the role of LIMS1, GCC2, as well as other non-HLA candidates paves the way for further detailed mechanistic studies with the potential for subsequent targeted therapeutics in case of an identified mismatch.

While the independent association of LIMS1 gene-level mismatches with DCGL was validated across both cohorts, intergenic regions were not evaluated in our approach – i.e., only loci within annotated gene bounds were considered. By integrating regulatory data from public datasets, we identified candidate SNPs in high LD with rs893403 or the LIMS1 haplotype, which could potentially represent otherwise causal cis-eQTLs for GCC2 (**Figure 7**). For instance, rs2460944 is highly linked (R^2^=0.96) with rs893403, has high RegulomeDB scores^30^, and is located within the transcription start site of GCC2. However, this and other candidates identified here will need experimental validation in future studies by altering these linked loci individually and evaluating GCC2 expression and clinical outcomes. We also acknowledge that we did not test for anti-LIMS1 antibodies in recipient sera, identified in recipients of high-risk mismatches in the prior report, which currently has no commercial assay.

In summary, using two prospective kidney transplant cohorts we identify a key role for D-R mismatches at the LIMS1 locus using a systematic screening approach of D-R mismatches at multiple genomic scales. Furthermore, our data reveals that these intronic variants at the LIMS1 locus have cis-regulatory function and D-R mismatches generated at these variants impact allograft outcomes without altering protein sequences. The novel mechanisms described here provide new insight into our understanding of D-R mismatches and are informative in ongoing efforts to improve long-term graft survival.

## Materials and Methods

### Genotyping and imputation for GoCAR and CTOT01/17

The genotyping, quality control (QC), and imputation for the GOCAR and CTOT cohort have been described in our previous study^21,32^. Briefly, donor DNA was obtained from either pre-perfusion allograft biopsies (in deceased donors) or PBMCs (in living donors), while recipient DNA was obtained from PBMCs. The extracted DNA was genotyped with Illumina Human OmniExpressExome Array for GoCAR and Illumina Infinium Global Screening Array for CTOT. Samples with: 1) genetically inferred gender not matched with reported gender; 2) missing genotype rate > 0.03; 3) excessive genome-wide heterozygosity, an indication of sample contamination were excluded. SNPs with: 1) missing rate > 0.05; 2) minor allele frequency (MAF) < 0.01; 3) Hardy-Weinberg equilibrium (HWE) p-value < 1e-6 were excluded.

We performed genome-wide genotype imputation on both cohorts. For GOCAR, the imputation analysis was done by the pipeline composed of SHAPIT^34^ and IMPUTE2^35^ software packages using the 1000 Genomes Project Phase I data^36^ as the reference panel; and for CTOT, the imputation was done by the Michigan Imputation Server (https://imputationserver.sph.umich.edu)^37^ using the Haplotype Reference Consortium (HRC) reference panel (Release 1.1)^38^. After imputation, an imputed genotype with a posterior probability <0.95 was set as missing data.

As shown in Figure S1, quality control on the imputed genome-wide genotype data of GoCAR and CTOT01/17 were performed separately in each cohort following the same strategy. The imputed SNPs with low confidence (INFO score < 0.4, GoCAR; r2 < 0.3, CTOT) or missing rate ≥ 5% were considered low quality and excluded. For instance, the rs893403 genotype was not confidently imputed in CTOT, resulting in 52/146 (35.6%) missing values in D-R mismatch calculation, and thus was not used for validation. SNPs with no alternative alleles across the samples (i.e., monomorphic) or within the MHC region (chr6:28866528-33775446 (hg19)) were excluded from downstream analysis as well. SNPs with mismatch carried by ≤ 5% of the D-R pairs were considered less frequent and removed from gene- and SNP-level screening analysis.

SNPs were annotated in terms of genomic locations (exonic, intronic, intergenic, etc.) or protein coding functions (synonymous, non-synonymous, frameshift, etc.) with annovar (version 2018Apr16)^39^ with Refseq hg19 assembly. The transmembrane or secreted genes were defined with the key words from Uniprot as described in the former study^5^, “Transmembrane [KW-0812]” OR annotation:(type:transmem) OR locations:(location:“Secreted [SL-0243]”) AND organism:“Homo sapiens (Human) [9606]”.

### Definition of D-R mismatch at different genomic scales

The D-R mismatch for each SNP was defined following the strategy in a former study^5^. A mismatch was defined as a donor carrying an allele that was not presented in the recipient. We consider mismatch derived from one “alien” allele introduced by donor as “single mismatch”, while two alleles introduced as “double mismatch”. The “single mismatch” and “double mismatch” were collapsed into a class named “any mismatch” (**Figure 1**). The mismatch score was calculated for each SNP among each donor-recipient pair. To define gene-level mismatch score, the mismatch status (0 for absence and 1 for presence) of all the SNPs within each annotated gene region were summed up. Similarly, to defnine mismatch score at different genomic scale, such as genome-wide, exonic, non-exonic, or all transmembrane or secreted gene regions, the mismatch status of all the SNPs within the corresponding genomic regions were summed as the raw score and further normalized by the inter-quartile range (IQR) of the corresponding raw scores across D-R pairs.

### eQTL and DEG analysis with DICE data

To explore the gene expression regulation effect of rs893403 and the identified haplotype in immune cells, we utilized the RNA-seq and genotype data generated from the DICE project (https://dice-database.org/). Request (request number: 97206-2) for the access to the DICE data deposited in the Genotypes and Phenotypes (dbGaP) database (accession number: phs001703) was approved. The detailed description of the dataset can be found in the original paper^18^. Briefly, whole transcriptome bulk RNA-seq was performed on 15 immune cell types isolated from leukapheresis samples of 91 healthy donors. Gene expression was measured as transcripts per million (TPM). The raw TPM expression profile was then log2-transformed by log2(TPM + 1). Genome-wide genotype data was generated by Illumina Infinium Multi-Ethnic Global-8 array, followed by imputation with the same pipeline as we applied to the CTOT cohort (see above). The number of risk alleles (A allele) of rs893403 and the number of minor alleles of the haplotype were counted in each sample. The association of expression of each gene with the genotype of rs893403 or the haplotype was tested with limma^40^, using an additive model of the number of risk alleles adjusted by age, gender, and race. Genes with nominal p-value ≤ 0.05 were identified as differentially expressed genes (DEGs). Gene set enrichment analysis of DEGs was performed with R package enrichR^41^, and gene sets with enrichment nominal p-value ≤ 0.05 were considered significant.

### Co-expression and functional enrichment analysis for GoCAR PBMC transcriptomes

The details of RNA-seq experiment and analysis on the PBMC of a subgroup of GOCAR patients at pretransplant and 3 months after transplant were described in our published studies ^14,42^. The normalized data was downloaded from the GEO database (accession number: GSE112927 and GSE120398). Gene co-expression was evaluated by Pearson correlation between the expression values of each gene pair for pre-transplant and 3-months post-transplant datasets separately. Genes with Benjamini-Hochberg adjusted p-value ≤ 0.05 and absolute correlation coefficient |R| ≥ 0.6 were considered co-expressed, followed by gene set enrichment analysis as described above.

### Online tools for in silico analyses

The linkage disequilibrium (LD) matrix (on GRCh37) for haplotype SNPs and rs893403 was generated with LDmatrix from Ldlink online portal (https://ldlink.nci.nih.gov/?tab=home) using all major continental populations from 1000 Genomics Project as reference. SNPs in LD with rs893403 and the haplotype were identified by LDproxy from LDlink online portal using the same reference. SNPs with R^2^>0.9 were considered in high LD with targeted SNPs, and each SNP was annotated with the Regulomedb database^28^ through LDproxy. The Regulomedb score was defined based on the strength of the supporting data as described in (https://regulomedb.org/regulome-help/), with lower value indicating stronger confidence of the regulatory role of the corresponding SNP. The SNPs were mapped to human kidney single cell ATAC-seq (scATACseq) peaks using Susztaklab Kidney Biobank^27^ with human genome hg19 assembly. The H3K27Ac mark, DNasel hypersensitivity clusters, transcription factor ChIP-seq clusters, chromatin state segmentation, and histone modification information were generated from UCSC Genome Browser with human genome hg19 assembly^43^. The eQTL analysis boxplot for LIMS1 gene expression associated with rs893403 and the haplotype () genotype in kidney tubulointerstitial tissue was generated from the NephQTL online portal (https://nephqtl.org)^17^. The LD structure of SNPs (on genome build hg19) within the LIMS1 gene region with the identified haplotype (represented by rs200106875) was evaluated by locuszoom/1.4^44^ with European population from the 1000 Genomes Project as reference.

### Statistical analysis

The associations of time-to-event outcomes (e.g., DCGL) with risk factors (e.g., D-R mismatches) were evaluated by Cox regression implemented in R package survival, with other relevant factors adjusted as covariates. Samples with missing data in covariates were omitted. For categorical- (e.g., rejection episodes) or ordinal-outcomes (e.g., CADI-, Ci-, Ct-, i-, t-scores at 12 months post-transplant), logistic- or ordinal regressions were performed to investigate the association with risk factors, respectively. Kaplan-Meier plot was generated by ggsurvplot implemented in R package survminer, and p-values for comparing between risk groups were generated by log-rank test.

## Acknowledgements

We thank all of the patients, the donors, and their families, the participating clinical sites, and the investigators in the GoCAR and CTOT-01/17 studies. MCM and ZZ acknowledge the Translational Collaborative Research Initiative Grant “Non-HLA donor-recipient differences and allograft survival,” provided by the Department of Medicine, and the computational resources and staff expertise provided by the Scientific Computing department at the Icahn School of Medicine at Mount Sinai.

MCM acknowledges funding from the American Heart Association (AHA) (15SDG25870018), the NIH (R01DK122164), and pilot funding from the CTOT-19 study (PSH, principal investigator; NIH grant U01AI063594) to study non-HLA D-R genetic differences. The data reported here are substudies of the GOCAR study (BM, principal investigator), supported by NIH grant U01AI070107-03, and the CTOT-01 study (PSH, principal investigator), supported by NIH grant U01AI63594-06. The content is solely the responsibility of the authors and does not necessarily represent the official views of the NIH. The authors sincerely thank the CTOT/GOCAR site investigators and staff for their efforts in collecting samples from the study participants.

Funding for the study “Impact of genetic polymorphisms on human immune cell gene expression” (DICE data) was provided by the NIAID (R24-AI108564). Data from the study were provided by Pandurangan Vijayanand on behalf of his collaborators at the La Jolla Institute for Allergy and Immunology.

