## Supplementary files for "Multiscale genetic architecture of donor-recipient differences reveals intronic LIMS1 locus mismatches associated with long-term renal transplant survival"

### Supplementary Figures

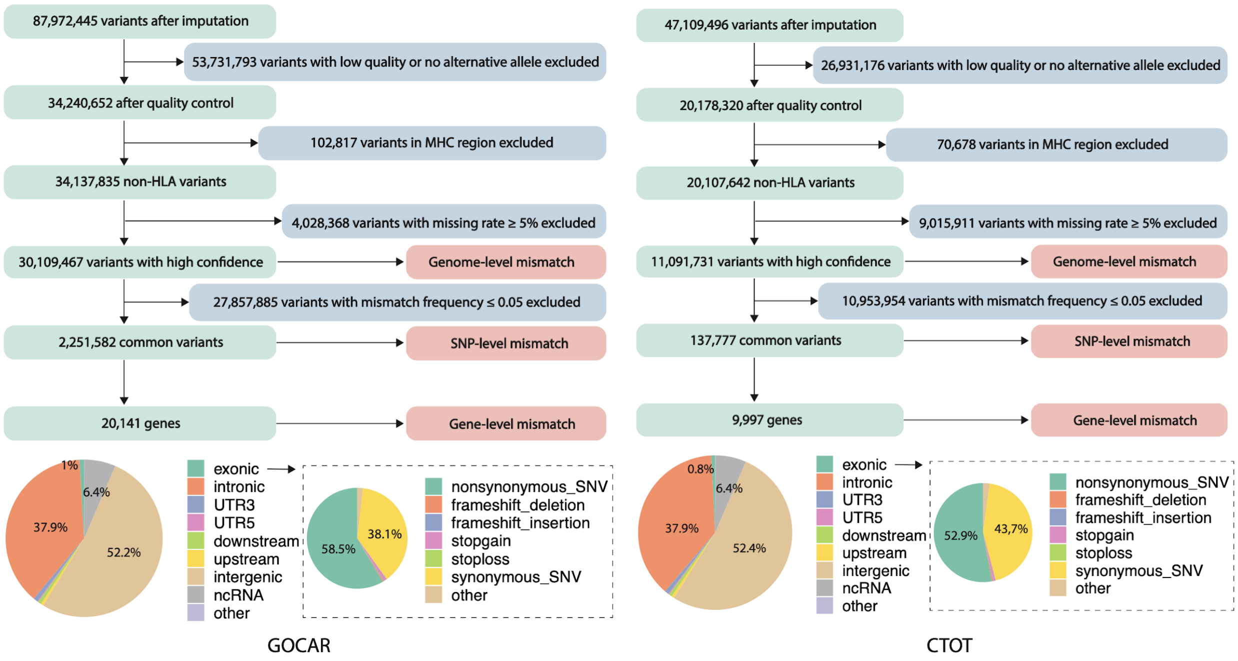


**Figure S1. Quality control of the imputed genome-wide genotype data for the GoCAR (left) and CTOT (right) cohorts.**

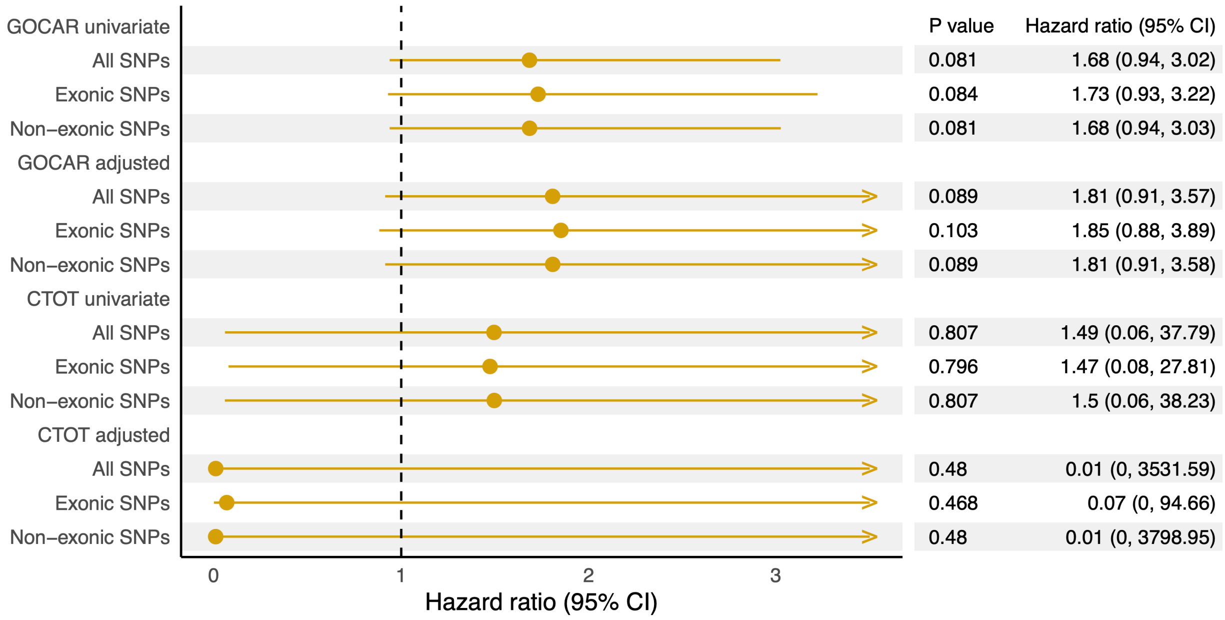


**Figure S2. Association of different genome-wide D-R mismatch scores with DCGL in European-to-European transplants in GoCAR and CTOT.** Genome-wide mismatch scores were calculated for all SNPs, exonic SNPs (SNPs located in exonic region), non-exonic SNPs (all SNPs minus exonic SNPs). Univariate and multivariable Cox regression (adjusting HLA mismatch score, induction therapy, and donor status) analyses were performed for both GoCAR and CTOT cohorts.

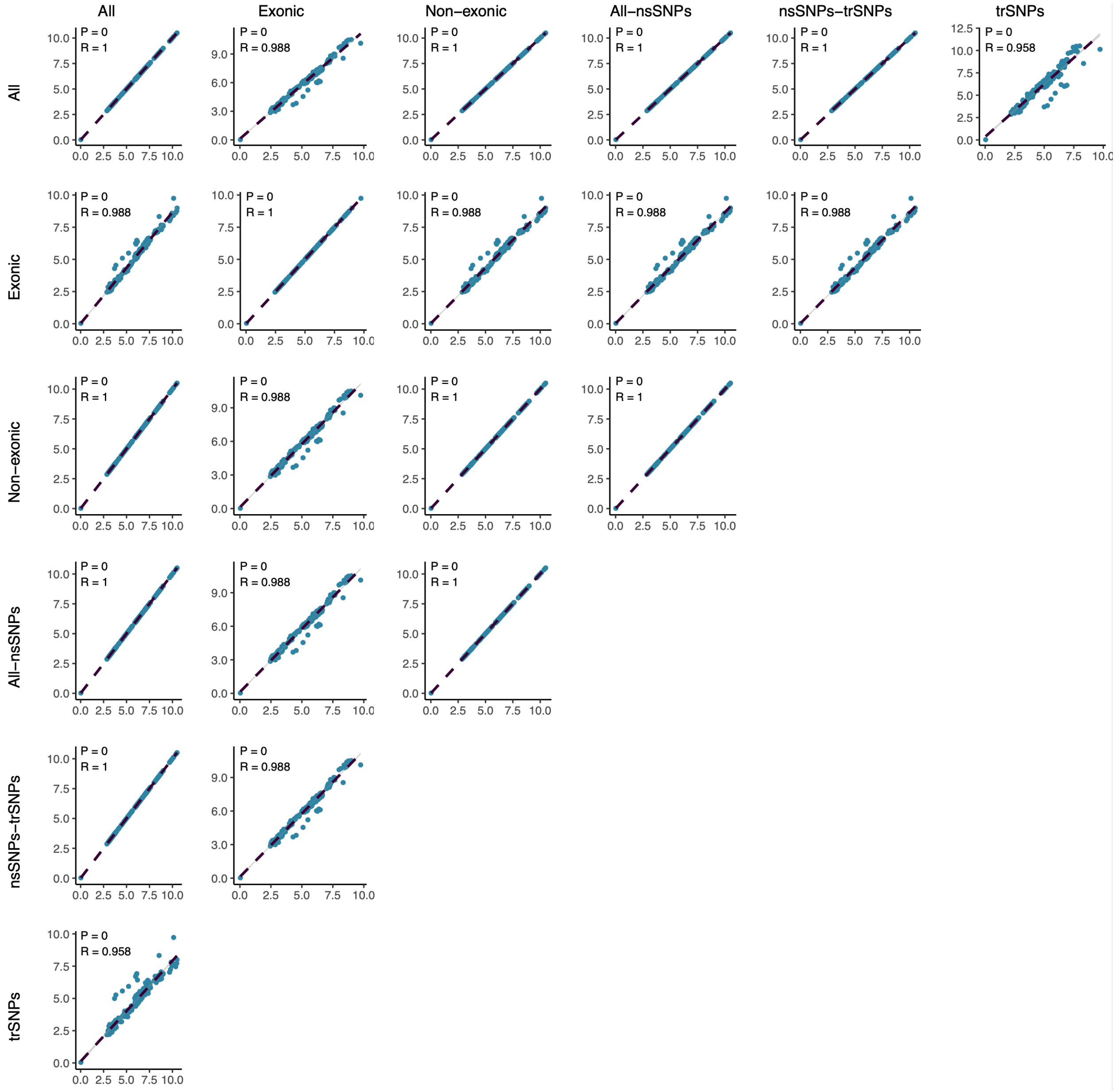


**Figure S3. Correlation among D-R mismatch scores defined within different genomic scopes in GoCAR**. SNP-level any mismatches were summed for all SNPs, exonic SNPs, non-exonic SNPs (all SNPs excluding exonic SNPs), all-nsSNPs (all SNPs excluding non-synonymous SNPs), nsSNPs-trSNPs (non-synonymous SNPs excluding transmembrane SNPs), and trSNPs (transmembrane SNPs) as raw counts, and normalized by their scope-specific IQR, respectively.

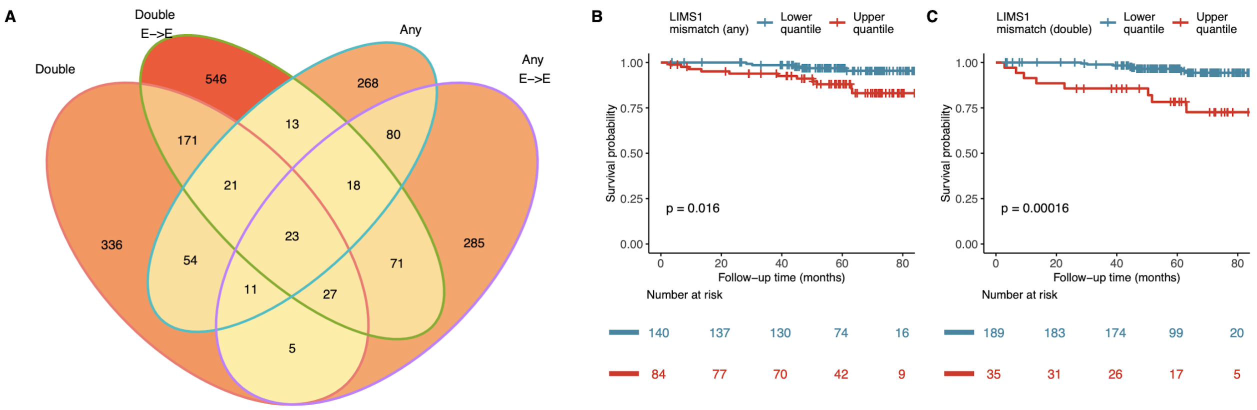


**Figure S4. Gene-level mismatches associated with graft loss.** (**A**) Venn diagram shows the number of genes identified with mismatch score significantly associated with DCGL (nominal p <= 0.05) from four different analyses: double mismatch or any mismatch (definition in Figure 1 and Methods) for the whole GoCAR cohort or the subset of European-to-European (E-to-E) D-R pairs. In GoCAR E-to-E D-R pairs, Kaplan-Meier plots show the graft survival curves for equally dichotomized groups of mismatch scores at LIMS1 locus, where mismatch scores were defined as “any mismatch” in (**B**) and “double mismatch” in (**C**). P-values were derived from log-rank tests in comparison of upper quantile versus lower quantile.

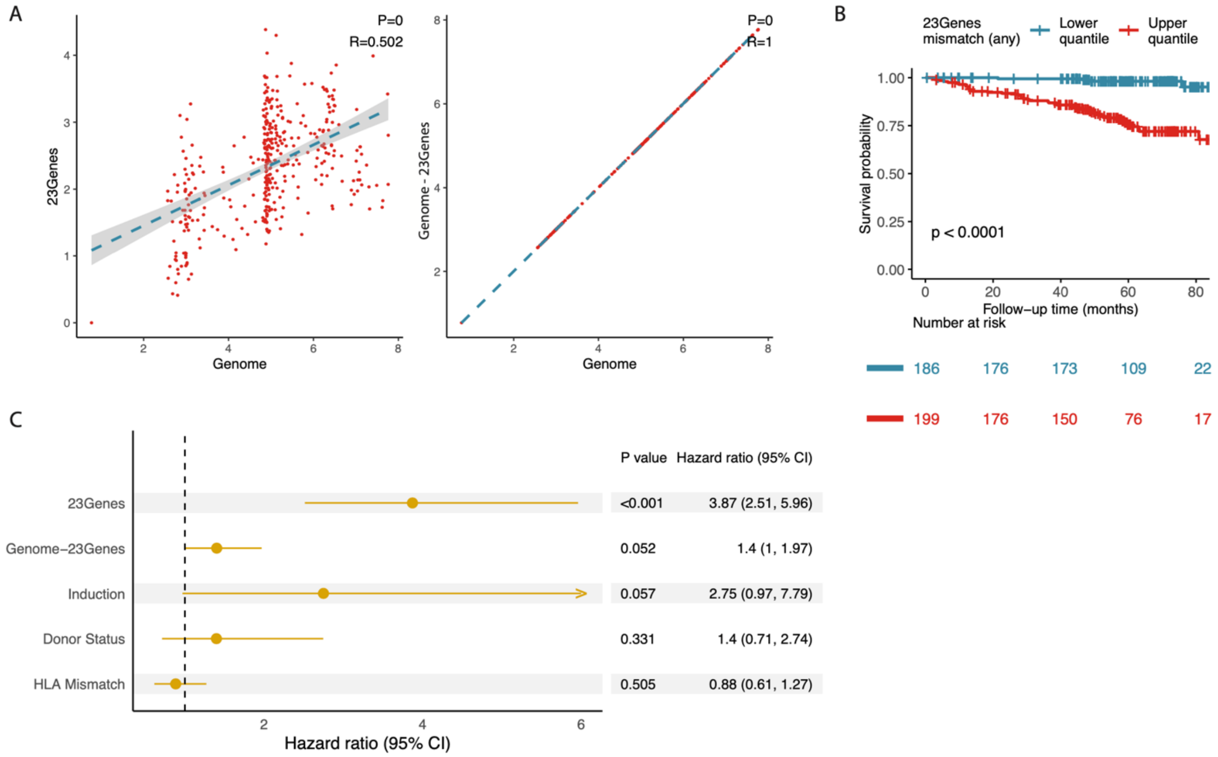


**Figure S5. Summary of mismatch score of the 23 candidate genes was significantly related to DCGL.** (A) The Correlation of the summary score of the 23 genes and other gene regions with genome-wide mismatch score. (B) Survival curve of the patients stratified by the 23 genes' mismatch score grouped by mean value. (C) Forrest plot of the hazard ratio of 23 genes' mismatch score to DCGL adjusted by mismatch score of the other genome regions, Induction therapy, donor status and HLA mismatch.

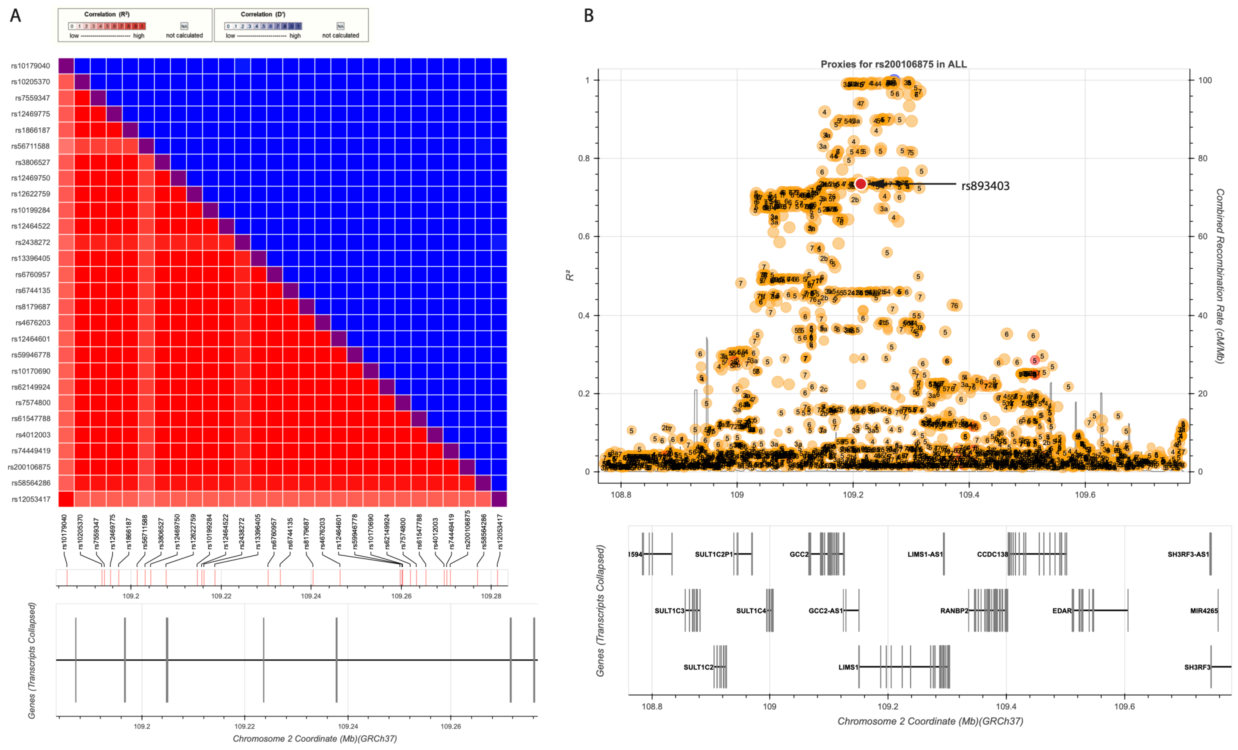


**Figure S6. Linkage disequilibrium between 30 top candidate SNPs and rs893403 within the LIMS1 locus.** (**A**) The LD metrics $R^2$ (red) and D' (blue) were calculated based on the genotype data from all the continental populations of the 1000 Genomes Project (hg19) using LDlink[1]. (**B**) The LD structure of the region surrounding the 30 candidate SNPs and rs893403 generated by LDlink, displayed as the $R^2$ value with a representative SNP rs200106875 of the haplotype of 30 candidate SNPs. The SNP rs893403 is highlighted by a red dot, clearly located in a distinct haplotype other than the 30-SNP haplotype. Gene models are shown below.

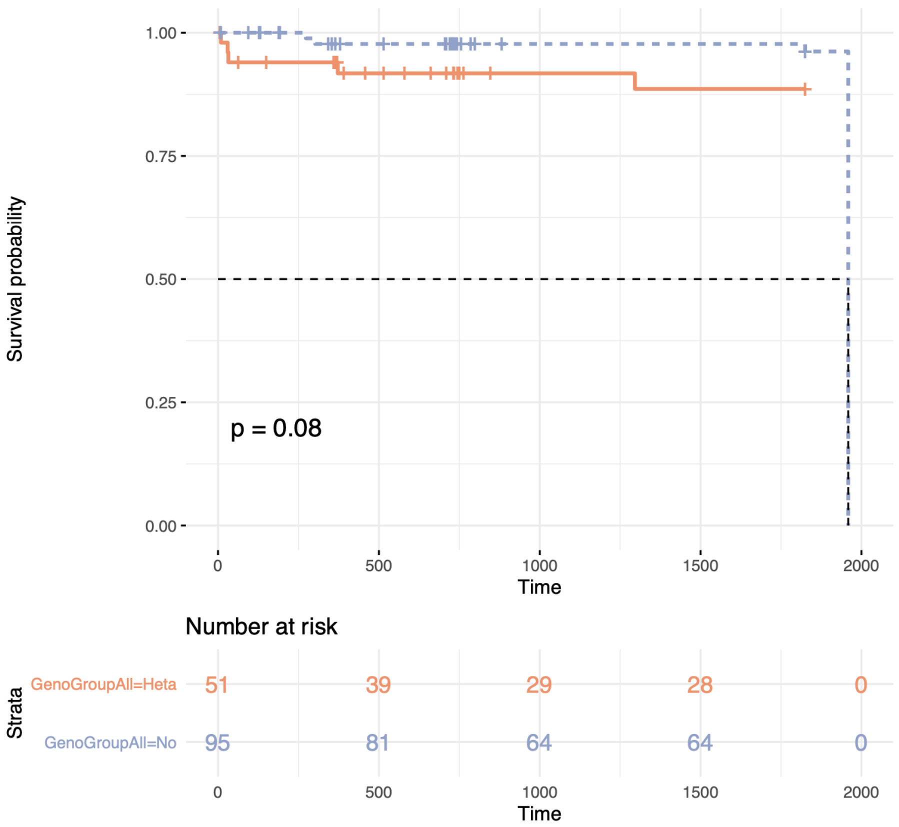


**Figure S7. Kaplan-Meier curves of DCGL for the CTOT patients grouped by presence and absence of any D-R mismatch of the identified LIMS1 haplotype.** P-value was derived from log-rank test.

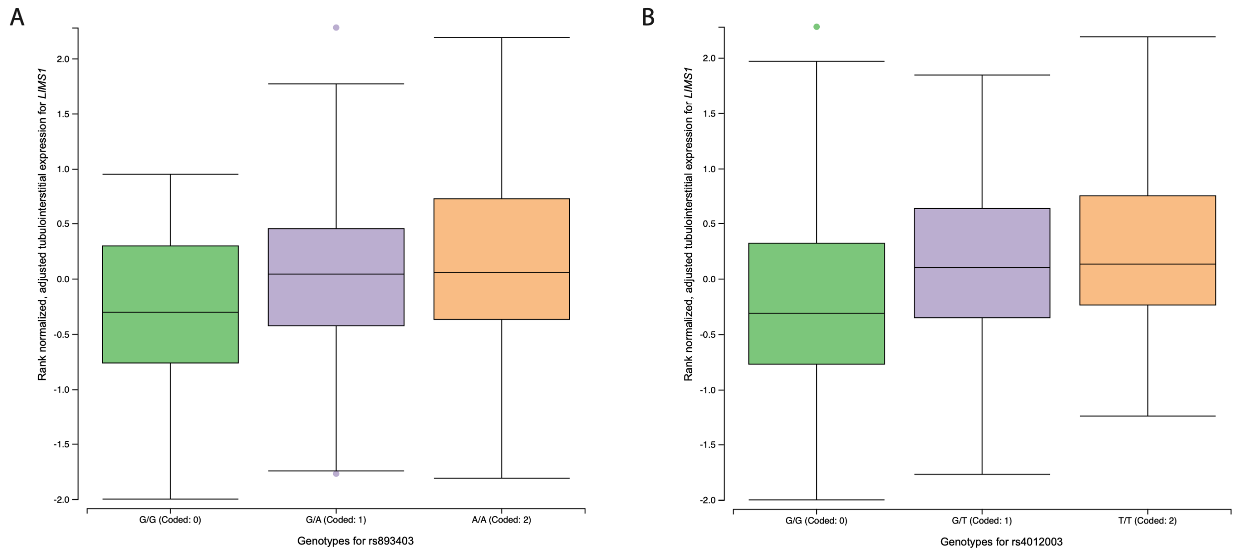


**Figure S8. eQTL data of rs893403 (A) and the haplotype (represented as one of the candidate SNP rs4012003) (B) in tubulointerstitial tissue from NephQTL[2].**

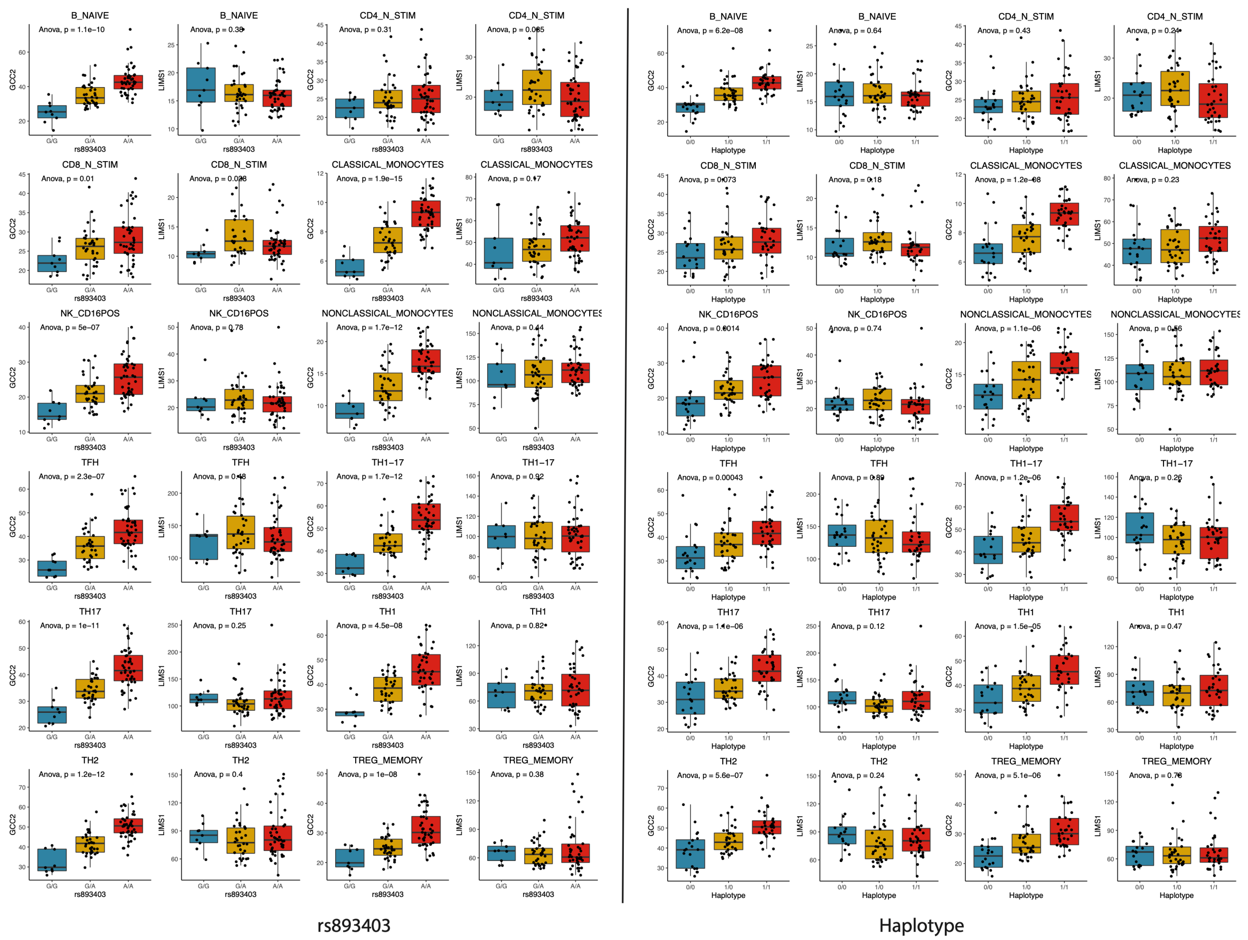


**Figure S9. eQTL analysis of GCC2 and LIMS1 using DICE data.** Box plots show the distribution of GCC2 and LIMS1 expression within each genotype group of rs893403 (left panel) and the haplotype (right panel) in 12 out of the 15 immune cell types from the DICE cohort (naive Treg, naive CD4$^{+}$ T cell and naive CD8$^{+}$ are shown in Figure 6A. The significance of the association between expression level and the genotype is indicated by the p-value derived from ANOVA.

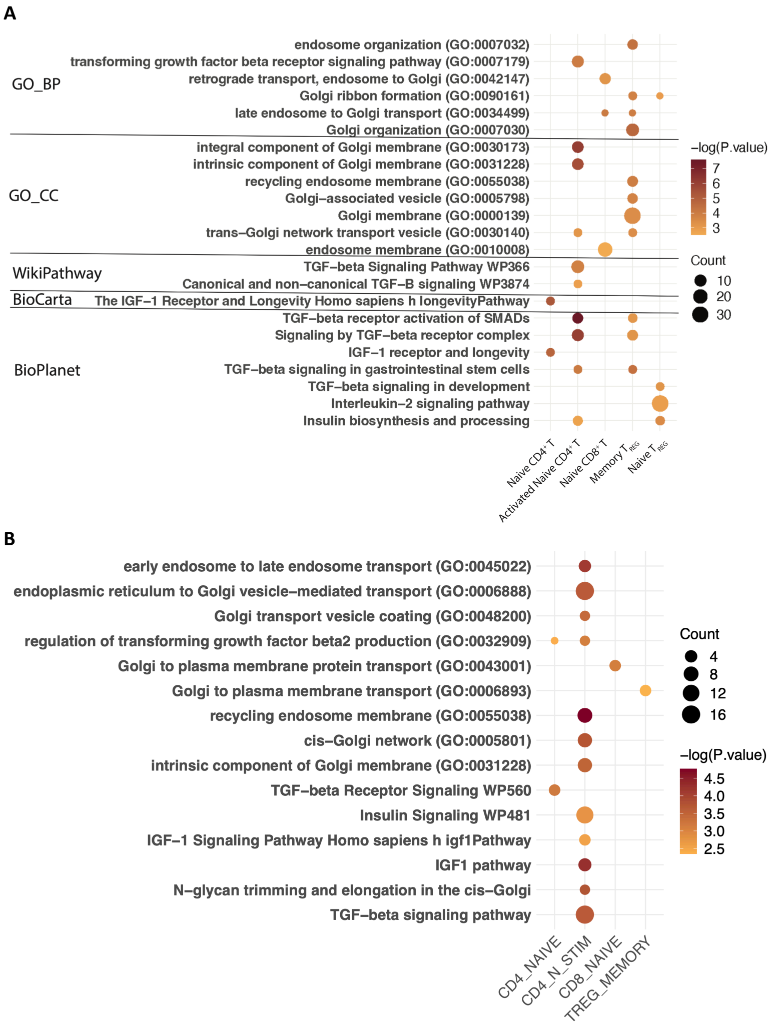


**Figure S10. Enriched functions of DEGs in relevant immune cell types from the DICE cohort.** Differentially expressed genes (DEGs) were identified by associating with the number of risk alleles of rs893403 (**A**) and the haplotype (**B**) with a nominal p ≤ 0.05.

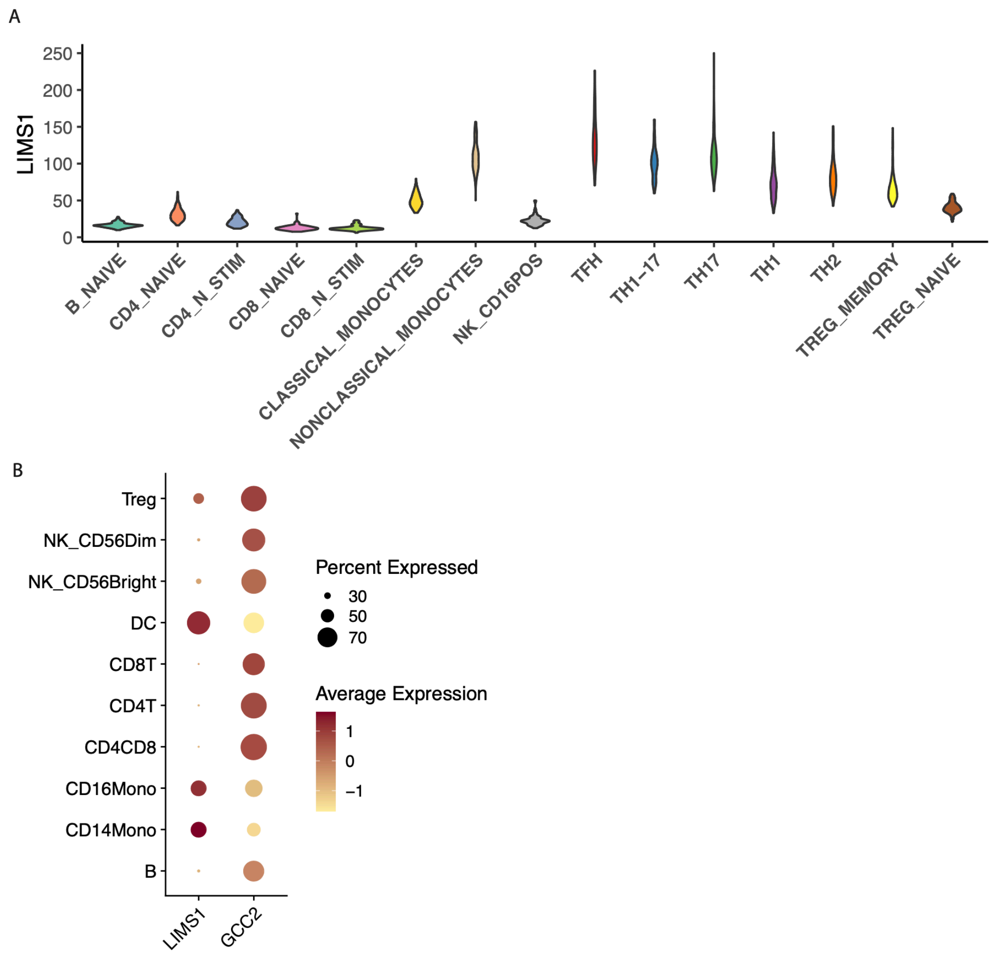


**Figure S11. Expression levels of LIMS1 and GCC2 in different immune cell types from published bulk and single-cell RNA-seq datasets.** (**A**) Distribution of LIMS1 expression in sorted PBMC subtypes from the DICE cohort. (**B**) Expression levels of LIMS1 and GCC2 in the PBMC single cell data.

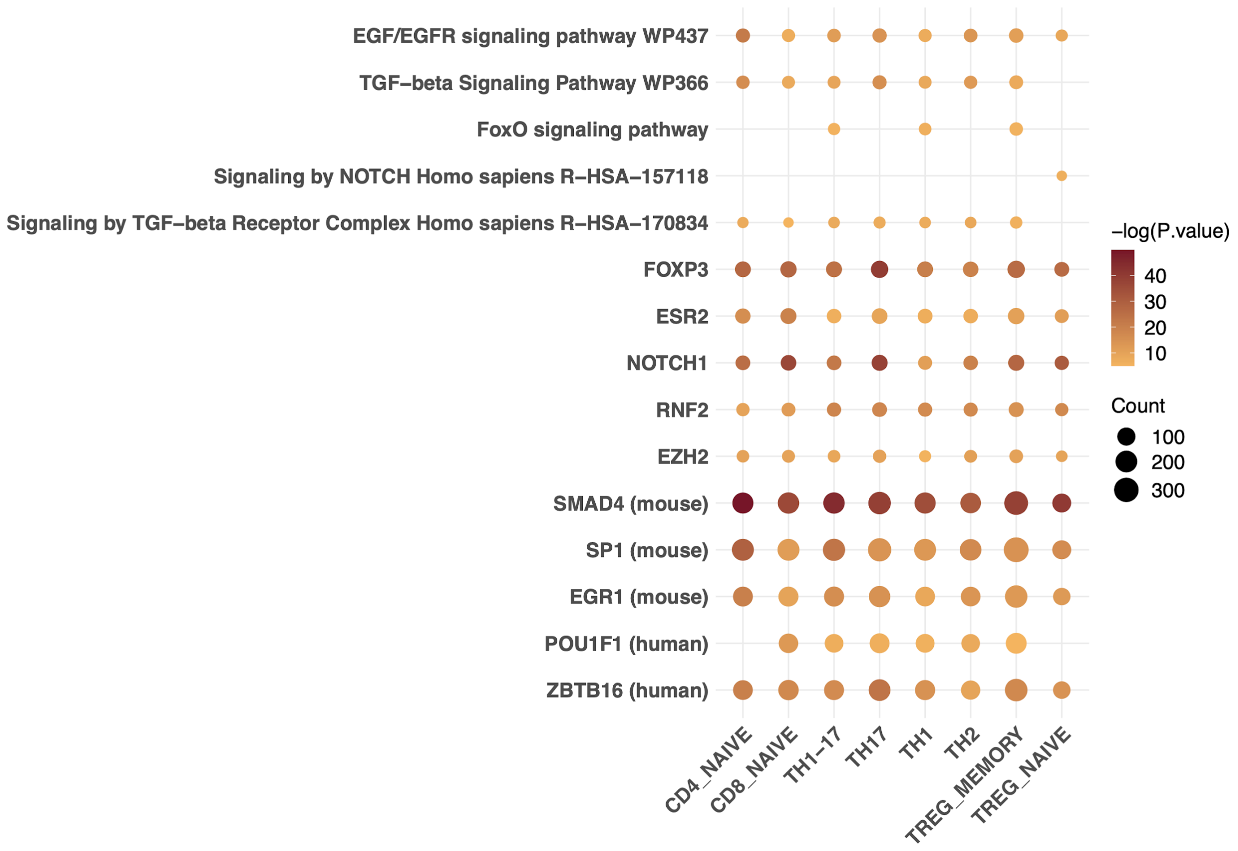


**Figure S12. Enriched functions and transcription factors of GCC2 co-expressed genes in multiple T cell subtypes from the DICE cohort.**

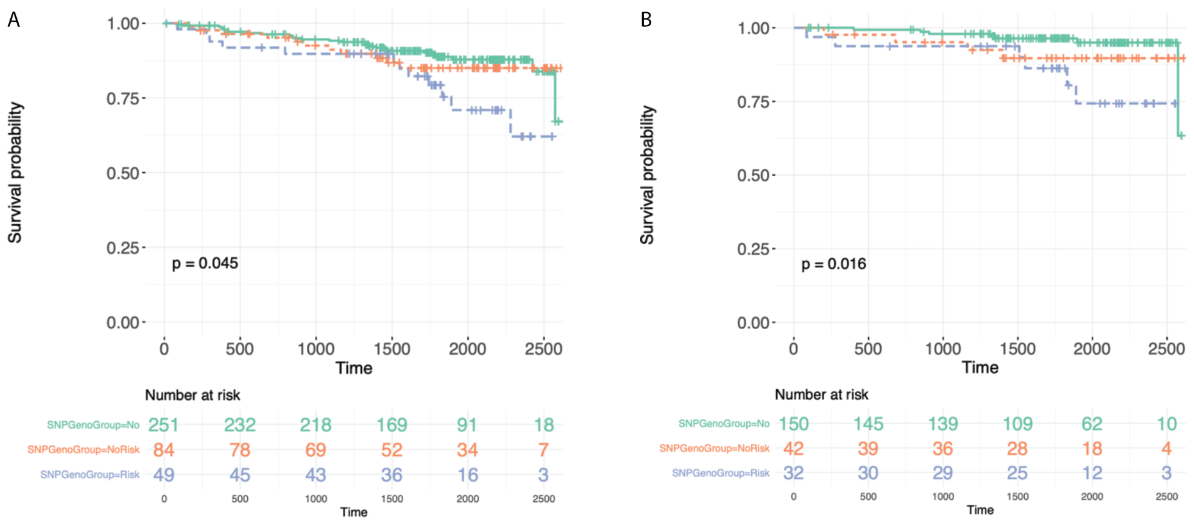


**Figure S13. Survival curves of DCGL in all patients(A) and E-to-E (B) of the GoCAR cohort grouped by rs2460944 risk allele (A allele) introduced by donor.**

### Supplementary tables

#### Table S1. Demographic and clinicopathologic characteristics of donors and recipients with genome-wide genotype data in the GoCAR and CTOT1/17 cohorts.

| Variable | GOCAR D-R pairs with genotype (n = 385) [a] | CTOT D-R pairs with genotype (n = 146) [b] | P-value[c] |
|---|---|---|---|
| **<u>Recipient</u>** | | | |
| **Death censored graft loss** (years) | | | |
| mean ± SD; median (range) | 4.6 ± 1.7; 4.9 (0.04, 7.3) | 3.7 ± 1.8; 5.0 (0.0, 5.4) | **<0.001** |
| # events (%) | 50 (13.0%) | 9 (6.2%) | **0.03** |
| **Age** (years), mean ± SD; median (range) | 49.9 ± 13.5; 50 (18, 83) | 43.5 ± 18.2; 47.5 (2, 89) | **<0.001** |
| **Gender**, male, n (%) | 257 (66.8%) | 88 (60.3%) | 0.19 |
| **Genetic ancestry**[d], n (%) | | | 0.40 |
| African American | 70 (18.2%) | 33 (22.6%) | |
| Asian | 13 (3.4%) | 2 (1.4%) | |
| Caucasian | 235 (61.0%) | 90 (61.6%) | |
| Hispanic | 67 (17.4%) | 21 (14.4%) | |
| **HLA mismatch score**[e], n (%) | 2.0 ± 1.0 | 2.8 ± 1.0 | **0.01**[f] |
| **Induction**, n (%) | | | 0.14 |
| No induction | 78 (20.3%) | 38 (26.0%) | |
| Non-depletional (IL2 antagonist) | 130 (33.8%) | 54 (37.0%) | |
| Depletional (Thymoglobulin or Campath) | 177 (46.0%) | 54 (37.0%) | |
| **<u>Donor</u>** | | | |
| **Age** (years), mean ± SD; median (range) | 42.6 ± 14.7; 45 (3, 73) | 39.3 ± 12.2; 38 (6, 65) | **0.01** |
| **Gender**, male, n (%) | 196 (50.9%) | 63 (43.2%) | 0.14 |
| **Genetic race**, n (%) | | | **0.006** |
| African American | 33 (8.6%) | 28 (19.2%) | |
| Asian | 7 (1.8%) | 2 (1.4%) | |
| Caucasian | 293 (76.1%) | 94 (64.4%) | |
| Hispanic | 52 (13.5%) | 22 (15.1%) | |

| | | | |
|---|---|---|---|
| **Donor type,** live donor, n (%) | 194 (50.4%) | 123 (84.8%) | **<0.001** |

[a]: Genome-wide genotype data is available for 385 donor-recipient (D-R) pairs from the parent GOCAR study after data processing and quality control detailed elsewhere[3].

[b]: Genome-wide genotype data is available for 146 donor-recipient (D-R) pairs from the parent CTOT study after data processing and quality control detailed elsewhere [ref KI].

[c]: P-value was calculated from unpaired t-test for continuous variables and from Fisher's exact test for categorical variables unless otherwise specified. Bold p-value < 0.05.

[d]: Genetic ancestry was inferred from genome-wide genotype data and considered more accurate than self-reported race[3].

[e]: HLA mismatch score was derived from 2-digit HLA allele typing. Following previous reports for GOCAR[3-5], the raw mismatch score (scaling from 0 to 6) was categorized into: 0 (no mismatches), 1 (1-2 mismatches), 2 (3-4 mismatches), and 3 (5-6 mismatches); while for the CTOT cohort, the raw mismatch score (scaling from 0 to 6) was used. In subsequent statistical analyses, this variable was used as numeric covariate in regression models.

[f]: In order to calculate the p-value, the raw HLA mismatch score used in CTOT was hereby categorized in the same way as GOCAR so that the HLA mismatch scores originally defined on different scales in the two cohorts are comparable. The p-value was calculated by Fisher's exact test.

**Table S2. Statistics of genome level mismatches between donor-recipient pairs.**

| | GoCAR | CTOT |
|---|---|---|
| Whole genome | 1,280,474.86 ± 335,138.36 | 233,365.3 ± 97,270.23 |
| Non-exonic SNPs | 1,272,112.08 ± 332,958.53 | 230,786.45 ± 96,425.89 |
| Exonic SNPs | 8,362.78 ± 2,205.68 | 2,578.85 ± 873.09 |
| Non-synonymous SNPs | 4,058.77 ± 1078.77 | 1,449.05 ± 468.12 |

**Table S3. Top candidate genes with gene-level D-R mismatches associated with DCGL in GoCAR.**

**Table S4. Association of LIMS1 mismatch with DCGL using univariate and multivariable Cox regression analysis in CTOT.**

| Variable | HR | 95% CI | P value |
|---|---|---|---|
| **Univariate analysis: D-R pairs of all ancestries** (n = 146; 9 [6.2%] graft loss events) | | | |
| LIMS1 gene level mismatch (any mismatch) (ref: no mismatch) | 4.08 | (1.21, 13.79) | **0.02** |
| **Multivariable analysis: D-R pairs of all ancestries** (n = 146; 9 [6.2%] graft loss events) | | | |
| LIMS1 gene level mismatch (any mismatch) (ref: no mismatch) | 6.08 | (1.46, 25.36) | **0.01** |
| Genome-wide mismatch | 1.73 | (0.66, 4.55) | 0.26 |
| Donor status (ref: Living donor) | 0.33 | (0.07, 1.50) | 0.15 |
| HLA mismatch score | 1.20 | (0.78, 1.83) | 0.41 |

**Table S5. Top candidate variants at the LIMS1 locus with D-R mismatches associated with graft loss.**

**Table S6. Association of identified LIMS1 haplotype with DCGL in CTOT using multivariable Cox model.**

| Variable[a] | HR | 95% CI | P-value |
|---|---|---|---|
| **D-R pairs of all ancestries** (n = 146; 9 [6.2%] graft loss events) | | | |
| Any haplotype mismatch (ref: no mismatch) | 4.79 | (1.03, 22.14) | **0.04** |
| Genome-wide mismatch | 1.87 | (0.93, 3.74) | 0.08 |
| Donor status (ref: Living donor) | 0.75 | (0.06, 1.30) | 0.11 |
| HLA mismatch score | 1.04 | (0.70, 1.55) | 0.83 |

[a]: Induction therapy was excluded from adjusted covariates because the model would have not converged when including the variable.

**Table S7. GCC2 expression was associated with SNP rs893403 and the haplotype genotype in multiple blood cell types from healthy individuals in the DICE data[6].**

**Table S8. List of SNPs in high LD with SNP rs893403 and located at peak regions of the LIMS1 locus in the kidney scATAC-seq data[7].**

**Table S9. List of SNPs in high LD with the haplotype and located at peak regions of the LIMS1 locus the kidney scATAC-seq data[7].**
